# Disentangling the Functional Roles of Pre-Stimulus Oscillations in Crossmodal Associative Memory Formation via Sensory Entrainment

**DOI:** 10.1101/2025.06.24.661289

**Authors:** Jan Ostrowski, Marike C. Maack, Michael Rose

## Abstract

The state of neural dynamics prior to the presentation of an external stimulus significantly influences its subsequent processing. This neural preparatory mechanism might be of particular importance for crossmodal memory formation. The integration of stimuli across different sensory modalities is a fundamental mechanism underlying the formation of episodic memories. However, the causal role of pre-stimulus neural activity in this process remains largely unclear. In this preregistered study, we investigate the direct relationship between transient brain states induced by sensory entrainment and crossmodal memory encoding. Participants (*n* = 105) received rhythmic visual stimuli at theta (5 Hz) or alpha (9 Hz) frequencies to evoke specific brain states. EEG recordings confirmed successful entrainment, with sustained increases in neural activity within the stimulated frequency bands persisting until stimulus onset. Notably, induced alpha oscillatory activity enhanced recognition memory performance reflected by increased sensitivity, and suggesting that alpha oscillations prepare the brain for optimal multisensory integration. These findings highlight the functional significance of distinct oscillatory brain states in facilitating memory encoding by increasing cortical excitability before stimulus presentation. Overall, our results emphasize the importance of pre-stimulus brain states in shaping the efficiency of memory formation across sensory modalities and shed light on how dynamic neural preparations support learning.

**Impact Statement:** By using sensory entrainment of pre-stimulus oscillations we could show thatalpha-band stimulation in particular enhanced crossmodal memory. These findings reveal a frequency-specific functional dissociation and highlight the potential of targeting preparatory brain rhythms to improve crossmodal memory formation.

## Introduction

Multisensory learning is fundamental for human cognition, enabling the encoding and retrieval of complex environmental information. In daily life, individuals continuously integrate sensory information from multiple modalities, such as visual and auditory stimuli, to enhance memory performance. This ability to form crossmodal associations supports essential cognitive functions, especially episodic memory formation (Dickerson & Eichenbaum, 2010; Gasser & Davachi, 2023). Given the relevance of multisensory learning, understanding the underlying neural mechanisms has become a key objective in cognitive neuroscience. Brain oscillations play a critical role in coordinating neural activity during multisensory learning. Theta oscillations (3–7 Hz) have been widely implicated in the formation of episodic memory, particularly in binding disparate elements of experience into coherent memories (Klimesch et al., 2011; Rudoler et al., 2023; Staudigl & Hanslmayr, 2013). Research suggests that theta rhythms support the temporal organization of information, facilitating associative encoding across modalities (Buzsáki & Moser, 2013; Herweg et al., 2020; Terada et al., 2017). Additionally, alpha oscillations (8–12 Hz) have been associated with attentional selection, serving as a gating mechanism to suppress irrelevant sensory input and enhance task-relevant processing (Foxe & Snyder, 2011; Jensen & Mazaheri, 2010; Waldhauser et al., 2012). However, the precise role of pre-stimulus theta and alpha dynamics of memory formation during multisensory learning remains unclear.

A growing body of evidence highlights the importance of pre-stimulus neural activity in shaping subsequent cognitive processing (Lindenbaum et al., 2023; Roberts et al., 2018; Salari & Rose, 2016; Taesler & Rose, 2022; Van Dijk et al., 2008; Zazio et al., 2022). Pre-stimulus theta and alpha power fluctuations have been linked to successful memory formation (Addante et al., 2011; Schneider & Rose, 2016; Scholz et al., 2017; Sweeney-Reed et al., 2016; Winterling et al., 2019), suggesting that oscillatory states before stimulus presentation may serve a preparatory function (Cruzat et al., 2021; Strunk & Duarte, 2019; Zoefel & VanRullen, 2017). In particular, we were able to support this notion in a previous investigation, where participants were required to memorize audiovisual pairs in a Subsequent Memory Effects task (SME; Ostrowski & Rose, 2024). We could demonstrate that theta and alpha oscillations have a significant impact on memory encoding during the pre-stimulus phase, as increases in theta (3–7 Hz) and alpha power (8–12 Hz) observed before stimulus presentation were associated with enhanced memory performance. Specifically, higher pre-stimulus theta and alpha activity has been linked to better recognition of crossmodal associations between stimuli, such as visual and auditory inputs.

These findings propose that pre-stimulus oscillations might optimize encoding conditions (Amil et al., 2024; Salari & Rose, 2016), aligning neural activity with upcoming information (Schneider & Rose, 2016; Terporten et al., 2019; Winterling et al., 2019; Yeh & Rose, 2019). However, a causal link between pre-stimulus oscillatory activity and successful learning has not yet been demonstrated. One promising approach is the modulation of pre-stimulus frequencies through entrainment. These methods, such as transcranial alternating current stimulation (tACS) and rhythmic sensory stimulation, provide the means to modulate oscillatory activity in a non-invasive manner (Bree et al., 2021; Neuling et al., 2015, 2017; Veniero et al., 2015). The application of external rhythmic stimulation can synchronize endogenous neural rhythms at targeted frequencies (Duecker et al., 2024; Notbohm et al., 2016; Notbohm & Herrmann, 2016; Thut et al., 2011), thereby affecting cognitive processes, and subsequently behavior (Bree et al., 2021; Michael et al., 2022; Wang et al., 2024). In sensory entrainment, neural oscillations are modified by an external visual or auditory stimulus during encoding. Depending on the sensory domain, either luminance or amplitude oscillate in a specific frequency, leading to increases in oscillatory power. As the brain synchronizes with these external rhythms, it may become more aligned at integrating sensory details into structured memories (Grover et al., 2021; Köster & Gruber, 2022; Singer, 1993; Wälti et al., 2020). Given the evidence that pre-stimulus oscillatory activity can affect memory performance, investigating whether externally applied rhythmic stimulation can modulate these oscillatory states to enhance learning is crucial to reveal a direct functional role of this neural mechanism. Furthermore, this might allow researchers to determine in a causal framework whether the potential enhancement of multisensory memory formation stems from improved temporal binding (theta) or more effective suppression of irrelevant information (alpha). However, studies investigating sensory entrainment in the context of multisensory learning have yielded mixed results so far (Hanslmayr et al., 2019; Wälti et al., 2020; Wang et al., 2018).

This pre-registered study aims to examine a direct link between pre-stimulus states of theta and alpha oscillations and multisensory memory formation by using visual sensory entrainment, while also addressing existing challenges in sensory entrainment through an optimized experimental paradigm. Using a between-subjects design, participants were required to memorize and later recognize pairs of visual and auditory stimuli. Visual sensory entrainment was presented immediately before each stimulus from the encoding task at either 5 Hz (theta group) or 9 Hz (alpha group). The choice of entrainment frequencies was based on observed effects from prior research where the same SME paradigm was used (Ostrowski & Rose, 2024). Arrhythmic stimulation was used as a control condition in which the entrainment oscillations were derived randomly from frequencies between 13 and 24 Hz. This approach extends previous work through a refined experimental design, allowing us to test whether pre-stimulus sensory entrainment might influence neural oscillations and memory performance.

Building upon prior research, the current study aims to replicate and extend previous findings through a refined experimental design that systematically manipulates brain oscillations before stimulus onset. First, we first expected that using an oscillating image as a stimulus for sensory entrainment will be successful in modifying oscillatory activity and hypothesized that it will lead to increased oscillatory power within the entrained frequency ranges (H1). Importantly, we hypothesized that both theta (H2a) and alpha (H2b) entrainment would enhance memory performance as compared to controls. Moreover, we hypothesized that theta and alpha entrainment might affect memory performance to a different degree, resulting in potential differences between the two conditions (H2c). Additionally, we expected that both theta and alpha entrainment might lead to improved memory performance as compared to no entrainment, which we assessed through a statistical comparison with the dataset from the previous study (H3).

## Results

In this study, participants (n = 105) performed in a sequential memory encoding and recognition task across three experimental runs, each containing audiovisual pairings that the participants were instructed to memorize **(Figure 1)**. Prior to each pairing in the encoding phase, participants were exposed to rhythmic visual stimulation at either theta (5 Hz) or alpha (9 Hz) frequencies, or exposed to arrhythmic stimulation (control) for two seconds. Each encoding run was followed by a short distractor task and a recognition phase, in which previously seen pairs were randomly intermixed with recombined lures. Participants indicated whether each pair was old or new via button press. We implemented an open-ended sequential design for gathering evidence, taking advantage of the Bayesian statistical framework. The data collection concluded either when at least moderate evidence had been gathered to accept or reject the null hypothesis for the respective contrast or when group size reached *k =* 35 for each group.

**Figure 1.**
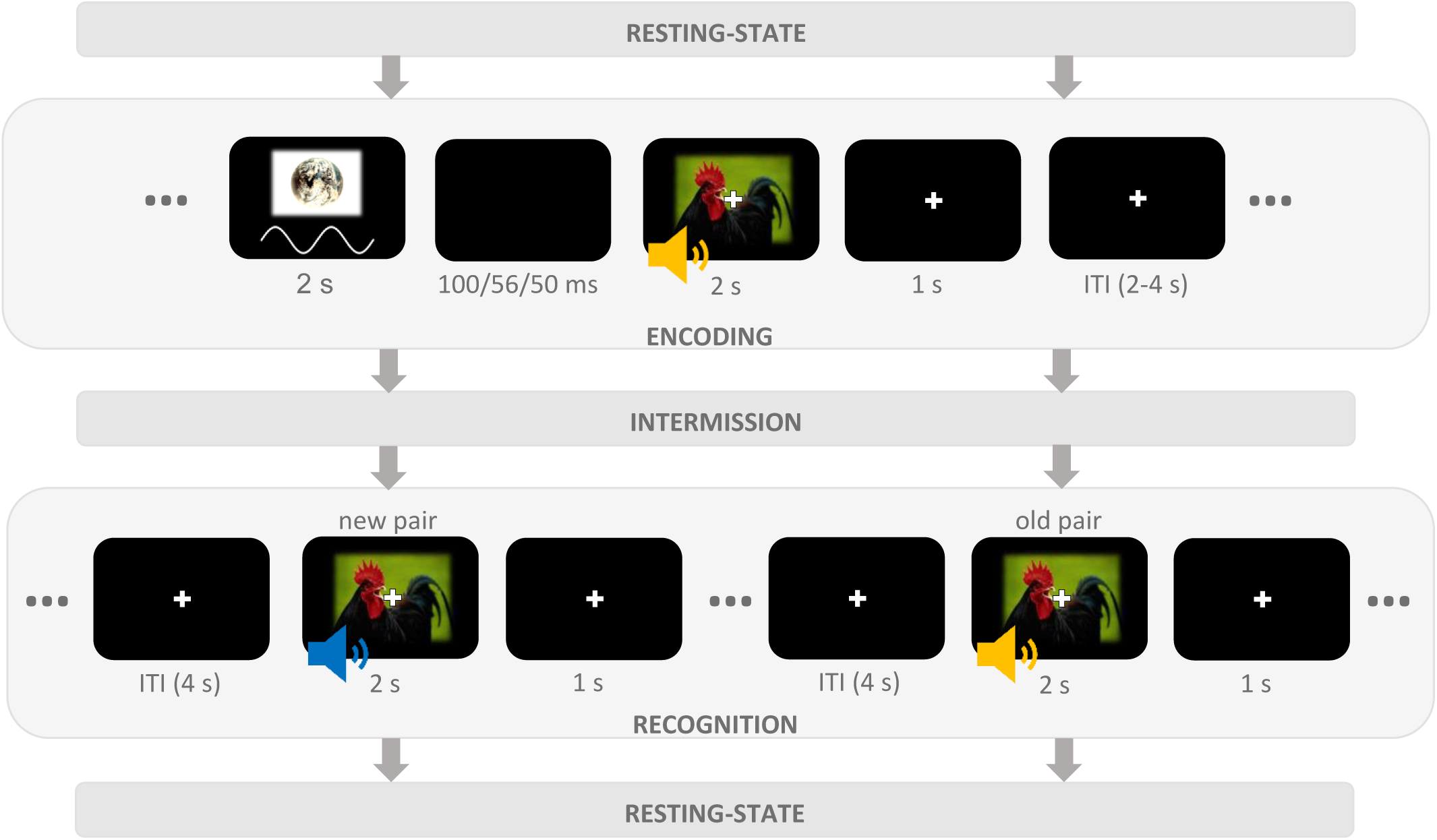
Schematic overview of one experimental run of the SME task. Each encoding trial began with a frequency-specific visual entrainment stimulus that lasted 2 s, followed by a black screen whose presentation duration differed depending on the group (theta: 100 ms; alpha: 56 ms; control: 50 ms). During entrainment, the luminance varied in a sinusoidal manner, resulting in a rhythmic oscillation of 5 Hz in the theta group and 9 Hz in the alpha group. Then, an image–sound pair was presented for 2 s, followed by a fixation cross for 1 s. Taken together, this comprised the possible window for responses. Participants judged whether both stimuli represented animals and were instructed to memorize each audiovisual combination. A fixation cross remained visible on the screen during the subsequent inter-trial interval of 2 to 4 s. In the recognition task, participants were presented with previously shown (old) and recombined (new) audiovisual pairs and indicated whether they remembered the particular combination of image and sound or not. Stimuli were shown for 2 s, and responses were recorded up to 3 s after stimulus onset. The inter-trial interval was fixed at 4 s, during which a fixation cross was shown. In the intermission between each encoding and recognition task, participants were presented with a short distraction task.

### Successful pre-stimulus visual stimulation modified targeted frequency

The focus of this study was to test whether visual sensory entrainment before the presentation of a stimulus would affect its subsequent encoding and thus result in changes in memory performance. As a prerequisite, we needed to make sure that the entrainment procedure would increase oscillatory power in the frequency bands corresponding to the entrainment frequencies (H1). To that end, oscillatory power in the late entrainment period (−1.1 s to –0.1 s relative to stimulus onset) from the theta and alpha entrainment groups was contrasted with the recorded activity from the control group. Two-tailed independent-samples *t*-tests were used on sample level, with a cluster-based permutation approach to account for multiple comparisons. The analysis was conducted for a frequency range of 1 to 40 Hz across the entire channel space. Comparing activity from the theta group with the control group, our analysis revealed a positive cluster ranging from 3 to 7 Hz and spanning the entire late entrainment period (*p* < .025, corrected, **Supplementary Figure S1A**), demonstrating the successful entrainment of pre-stimulus theta activity. Simultaneously, a negative cluster was observed that ranged from 13 Hz to 40 Hz (*p* < .025, corrected), covering most of the beta as well as lower gamma bands. In the comparison between the alpha group and control group, a positive cluster was found in the range of 6 to 10 Hz that spanned the whole analysis window (*p* < .025, corrected), also showing the specific entrainment of alpha band oscillations before the onset of the stimulus pair. Furthermore, the analysis revealed a negative cluster in the high beta/low gamma band ranging from 29 to 34 Hz, spanning the whole analysis window as well (*p* < .025, corrected). Generally, the entrainment seemed to be centered around occipital and parieto-occipital electrodes, and the effects in all entrainment groups were observed only in the pre-stimulus period (**Figure 2**), since our analysis revealed that oscillatory activity after stimulus onset did not differ between the theta, alpha, and control groups (*p* = .069; **Supplementary Figure S2**) In addition, comparing pre-stimulus power from both entrainment groups with oscillatory activity from the NE group using identical analysis parameters revealed similar patterns. Specifically, we found a significant positive cluster in the theta frequency range (theta vs NE; *p* < .025, corrected) as well as in the alpha band (alpha vs NE; *p* < .025, corrected). The common effects found in the control as well as in the NE contrast are shown in **Figure 2B** and **2C**, demonstrating the specificity of the different entrainment protocols (for visualizations of individual contrasts, see **Supplementary S2**). These results suggest that the entrainment of 5 Hz in the theta group and 9 Hz in the alpha group successfully modified oscillatory activity in the pre-stimulus window and the targeted frequency selectively and consistently. As no alpha modification was observed in the theta group, and no theta modification in the alpha group, the observations support our hypothesis that sensory visual entrainment can selectively modify ongoing oscillations (H1).

**Fig. 2.**
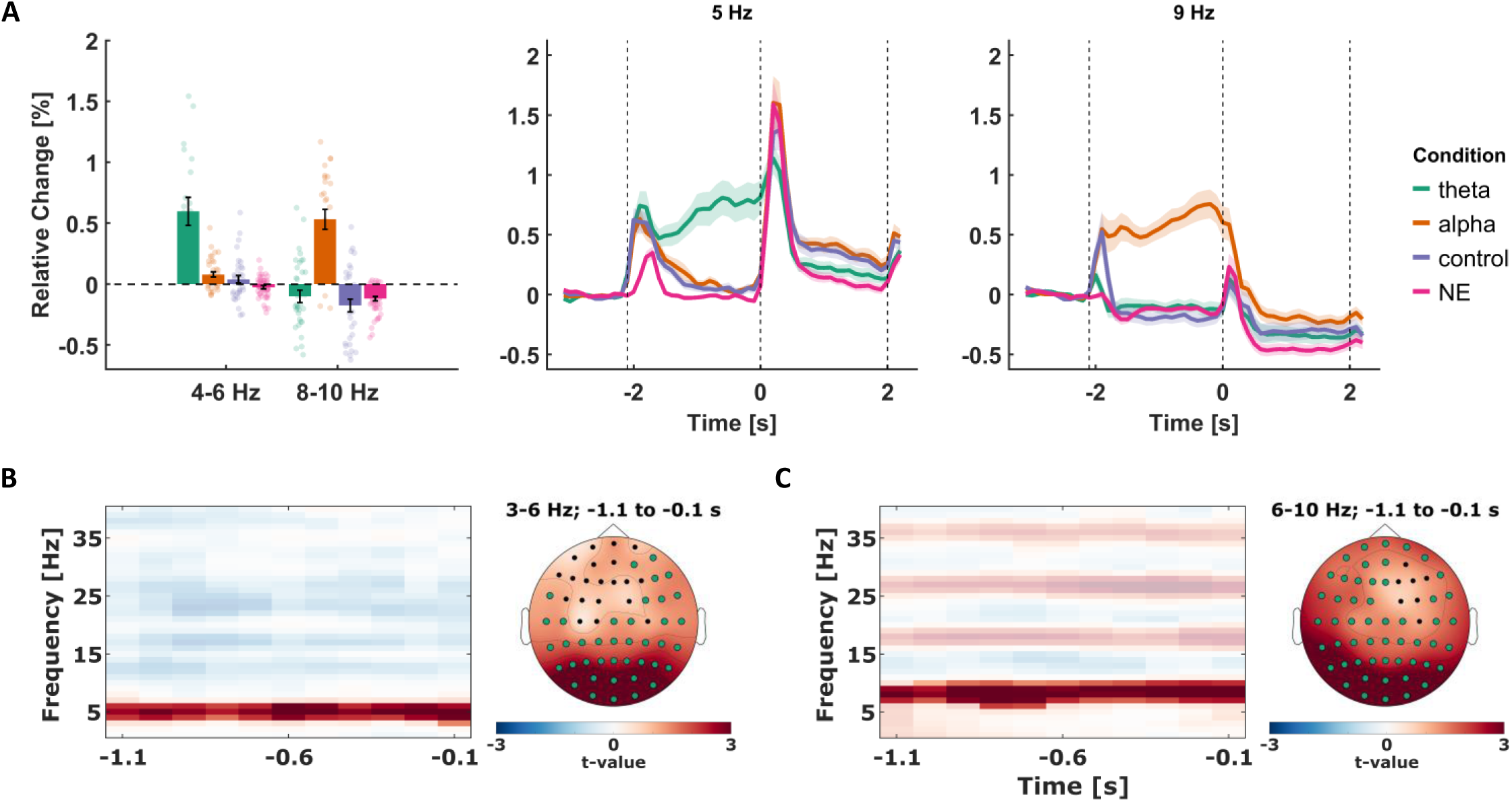
Effects of visual sensory entrainment on recorded EEG activity. **(A)** *Left* shows the average relative change of oscillatory power in reference to the baseline period for every group. Specifically, the average change is displayed for the mean activity from the 5 Hz and 9 Hz envelopes (± 1 Hz). Each data point shows the respective mean value for one participant, and the black error bars represent the standard error of means. *Right* shows the average time courses of relative change in power for the 5 Hz and 9 Hz narrow bands across the whole trial period for every group. The shadings around the lines represent the standard error of means across participants. All three figures depict relative change derived from the mean of the occipital electrodes *O1, O2,* and *Oz*. The first dashed vertical line marks the onset of the entrainment stimulus, while the other two mark the stimulus presentation window. **(B)** Visualization of the common effects of visual entrainment that were found in the contrast of the theta group with the control group, as well as in the contrast with the NE group. The time-frequency plot (*left*) shows the dimension of the common cluster along the time and frequency dimensions, depicting the average *t-*values across all contributing electrodes. Positive *t*-values signify greater relative change in the theta group, while opaque data points mark a significant difference at *p* < .025 (corrected). On the *right*, the topographical distribution of the common effects is shown, with electrodes contributing to the cluster marked in green. **(C)** Same as in (B) but for the alpha entrainment group.

In some studies, the entrainment frequency is individually tailored to match participants’ endogenous rhythms (Duecker et al., 2024; Zaehle et al., 2010). This approach is particularly common in alpha entrainment research, where resting-state EEG is used to identify the individual alpha frequency (IAF) as a target for stimulation (Janssens et al., 2022; Kasten et al., 2019; Klimesch, 2012; Stecher et al., 2017). To explore whether the match between stimulation frequency and endogenous alpha rhythms modulated entrainment strength in the present study, we computed the absolute difference between each participant’s IAF and the stimulation frequency in the alpha group (9 Hz). The IAF was extracted from resting-state EEG recorded prior to the main experiment by calculating power spectra using a multitaper fast Fourier transform (1–40 Hz). It was defined as the frequency showing the maximum power within the 8–12 Hz range, averaged across posterior electrodes (Pz, POz, Oz, O1, O2). Thus, we correlated this IAF distance with the maximal relative change in alpha power during the recognition phase, as an index of entrainment strength. This analysis was restricted to participants in the alpha entrainment condition. The correlation was not statistically significant, *r*(43) = –0.034, *p* = .849, indicating that the individual distance from the stimulation frequency did not predict the strength of neural entrainment as indexed by maximal alpha power modulation. Surprisingly, the same analysis for theta revealed a significant negative correlation between the individual theta frequency (ITF) distance and the maximal relative change in theta power during the encoding phase, *r*(55) = –0.296, *p* = 0.028. This suggests that a smaller difference between the individual theta frequency and the entrainment frequency might be associated with greater increases in theta power.

### Alpha but not theta entrainment enhances memory performance

In this study, we aimed to determine whether changes in oscillatory activity during the pre-stimulus interval could causally influence an individual’s ability to encode audiovisual associations. First, the analysis of performance in the categorization task during encoding yielded moderate evidence in support of the null hypothesis, BF₁₀ = 0.381, suggesting no significant differences in accuracy across conditions. These findings suggest that participants consistently adhered to task demands throughout the experiment, supporting the validity of subsequent analyses on oscillatory activity and memory performance (see **Supplementary 3** for further details). Notably, memory performance in the recognition task as measured by the sensitivity index *d′* was significantly enhanced in the alpha entrainment group (*M* = 1.46, *SD* = 0.60) as compared to the control group (*M* = 1.18, *SD* = 0.50; **Figure 3A**). An independent-samples *t*-test yielded a Bayes factor of BF₁₀ = 3.29, providing moderate evidence for the alternative hypothesis and suggesting that increased alpha-band activity induced by visual entrainment may facilitate the formation of audiovisual associations (H2b). Further analysis revealed that this effect was primarily driven by a measurable increase in hit rate in the alpha group (M = 61.498%, SD = 14.443%) as compared to the control group (M = 49.792%, SD = 15.029%), BF₁₀ = 22.5742. Simultaneously, no differences were observed in the false positive rate between the groups (alpha: M = 13.9%, SD = 6.67%; control: M = 13%, SD = 5.73%), BF₁₀ = 0.2852 (**Figure 3B**). This suggests that participants in the alpha group were more likely to correctly recognize an old stimulus pair compared to those in the control condition. In contrast, the comparison of sensitivity between the theta group (*M* = 1.28, *SD* = 0.55) and the control group yielded a Bayes factor of *BF*_10_ = 0.49, indicating weak evidence for the null-hypothesis. Similarly, the direct comparison between the theta and alpha groups resulted in a Bayes factor of BF₁₀ = 0.53, further suggesting weak support for the null hypothesis. Although these results do not support our hypotheses H2a and H2c, they provide evidence that any effect of theta entrainment on encoding performance may be smaller or more variable than anticipated. Together, these findings point to a potentially specific role of alpha oscillations in enhancing audiovisual memory encoding, highlighting the importance of frequency-specific mechanisms in pre-stimulus neural dynamics.

**Figure 3.**
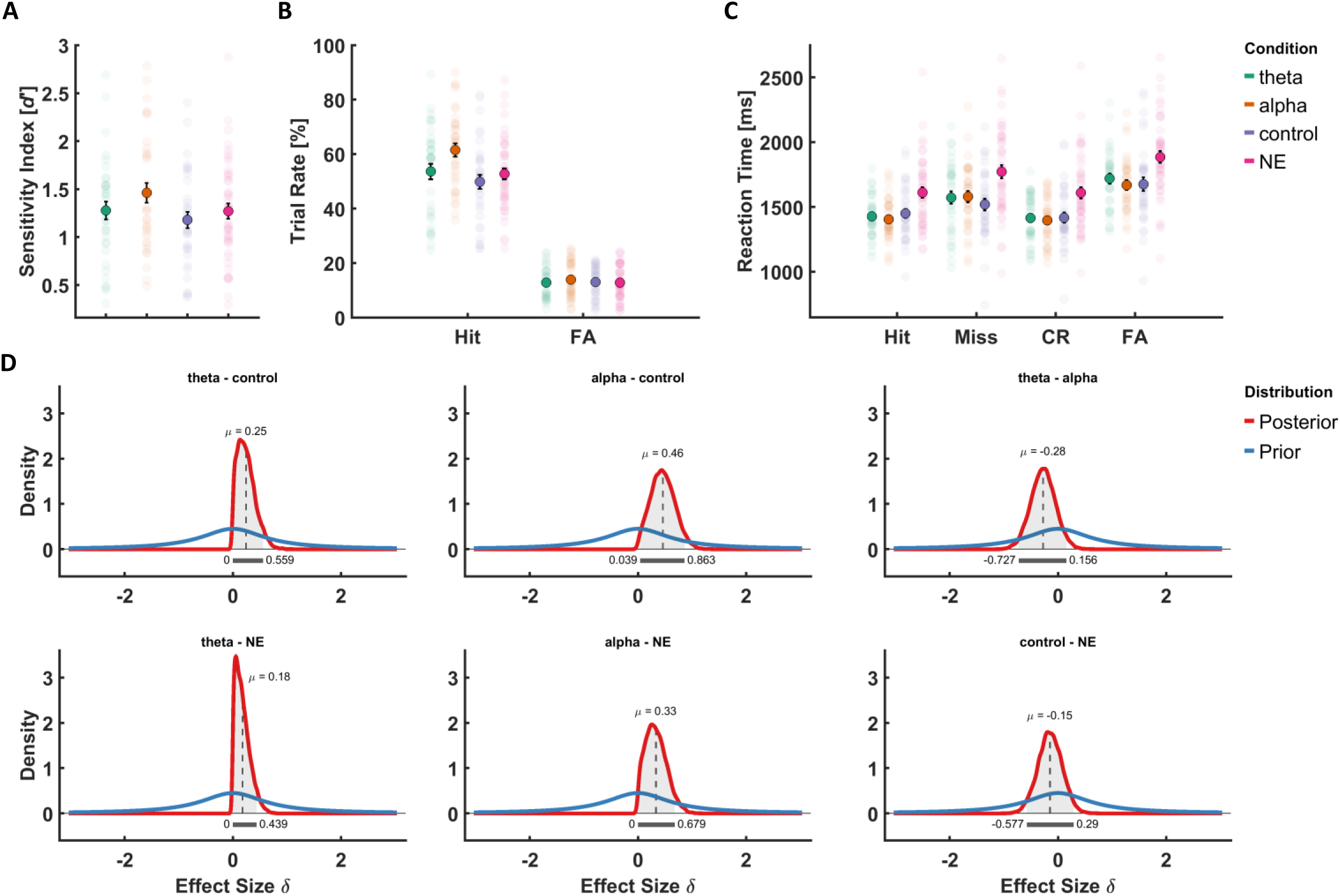
Effects of neural entrainment on recognition memory performance. **(A)** Mean sensitivity index (*d’*) with standard errors of the mean (SEM) across the three entrainment conditions **(B)** Average trial rates depicting the proportion of remembered old pairs (Hit) and new pairs erroneously categorized as old (false alarms; FA) across individuals for each group. Black error bars mark the SEM. (**C)** Group means of reaction times (RTs) for all response categories from the recognition task illustrating differences in processing speed between the groups. In addition to Hits and FAs, the figure includes RTs for not recognized old pairs (Miss) and correctly rejected new pairs (CR). Individual group means are reported in **Supplementary 5**. **(D)** Prior and posterior distributions for the individual group contrasts for sensitivity indices. The Highest Density Interval (HDI) is marked by the grey bar and the light grey shaded area under the curve of the posterior distribution. The mean effect size *μ* is marked by the dashed line.

To address our preregistered hypothesis (H3), we compared memory performance in the entrainment groups to that of participants from a previous study who were not exposed to any rhythmic stimulation during the pre-stimulus interval but instead viewed a static fixation cross (NE group; M = 1.27, SD = 0.53). The results of the Bayesian *t*-test showed that neither the theta group nor the control group differed significantly from the NE group, BF₁₀ = 0.26 and BF₁₀ = 0.16, respectively, indicating moderate-to-strong evidence for the absence of an effect. The comparison between the alpha group and the NE group yielded a Bayes factor of BF₁₀ = 1.03, indicating that the data did not provide conclusive evidence for either hypothesis. However, we found moderate evidence suggesting that the hit rate measured in the alpha group still differed from the hit rate in the NE group (M = 52.662%, SD = 13.724%), BF₁₀ = 3.8563. Again, we also found moderate evidence that the false positive rate from the NE group (M = 12.8%, SD = 6.1%) was likely not statistically different from the false positive rate in the alpha group. This indicates that participants receiving alpha band stimulation correctly remembered old stimuli more often than participants who were not stimulated at all. In addition, we investigated whether sensory entrainment might have affected how memory performance changed across the experiment. While we found that memory performance generally increased over the course of the experiment, this effect was not modulated by the pre-stimulus condition, BF = 0.0343, indicating that the improvement was consistent across entrainment conditions (see **Supplementary 4** for further details).

### Visual entrainment accelerates response times

Next, we explored the effect of visual entrainment on response times. First, we compared group means of reaction times from the categorization task during encoding using the Bayesian version of a one-way ANOVA with the factor *pre-stimulus condition* (theta, alpha, control, NE). The analysis revealed moderate evidence in favor of the alternative hypothesis, BF_10_ = 5.0737, suggesting relevant differences in response times between the groups during encoding. Further analysis revealed that participants in the NE group responded significantly slower as compared to the entrainment groups (see **Supplementary 3** for details). Next, differences in reaction times from the recognition task were assessed using the same statistical approach but conducted separately for all four response categories (hits, misses, correct rejections, false alarms). Importantly, we found strong evidence that response times differed significantly between conditions for every response category (hit: BF_10_ = 78.0847; miss: BF_10_ = 8.0419; correct rejections: BF_10_ = 49.3906; false alarms: BF_10_ = 18.2397). Subsequent analyses revealed that the entrainment groups did not differ in response times, regardless of which response category was tested. The NE group, however, displayed consistently slower reaction times than the other groups for every response category (**Figure 3C**; see **Supplementary 5** for more information on the individual group contrasts). The results suggest that participants who received visual stimulation during the encoding phase responded faster in the recognition phase than participants who were not stimulated. In addition, we investigated whether the entrainment condition would affect the discrepancy in response times between recognition trials with correct and incorrect responses. To that end, response times from all response categories were grouped according to the correctness of the corresponding trial and submitted to a Bayesian mixed-design ANOVA. The analysis yielded strong evidence in favor of the null hypothesis, BF_10_ = 0.1161, indicating that the difference in response times between correct and incorrect trials was not modulated by the entrainment condition.

### No lasting effect of entrainment condition on resting state activity and salience reports

To complement our pre-registered analysis, we conducted exploratory investigations examining resting-state EEG data before and after the experiment, as well as differences between groups in subjective salience reports. Comparing the difference in resting-state activity from before and after the experiment across the theta, alpha and control groups revealed no notable differences (p = .2972, corrected). This suggests that any changes in baseline activity due to the experiment were not dependent on the entrainment condition and appeared to be a general effect instead (see also **Supplementary 6**). To control for the subjective experience of the visual entrainment, participants rated its pleasantness and salience as well as their own perceived attention and fatigue at the end of every encoding task. Although we observed noticeable decreases in perceived attention as well as increases in perceived fatigue over the course of the experiment, BF_attention_ = 1.855 x 10^9^, BF_fatigue_ = 0.3415 x 10^15^, these effects did not interact with the entrainment condition, BF_attention_ = 0.195, BF_fatigue_ = 0.0552 (**Supplementary 7**). Overall, these results suggest that subjective perceptions and task engagement were comparable across groups, reducing the likelihood of confounds influencing the behavioral outcomes.

## Discussion

This study aimed to investigate the causal role of pre-stimulus oscillations in the encoding of crossmodal associations. Using visual sensory entrainment targeted at theta (5 Hz) and alpha (9 Hz) frequencies, we aimed to modulate neural rhythmic activity prior to stimulus presentation and assess its impact on memory performance. Our main findings demonstrated successful frequency-specific entrainment of pre-stimulus oscillations. Notably, alpha-band entrainment before stimulus presentation resulted in significantly improved recognition memory, as evidenced by increased sensitivity driven by higher hit rates. In contrast, pre-stimulus theta entrainment did not produce measurable behavioral effects. These results support a functional dissociation of pre-stimulus alpha and theta oscillations in relation to memory encoding, with alpha activity playing a more prominent role in facilitating successful associative memory formation. Importantly, these effects were driven solely by transient pre-stimulus modulation, with no evidence for lasting entrainment effects during stimulus presentation or changes in post-experiment resting-state activity, enabling the differentiation of their respective functional contributions.

Previous research has demonstrated that pre-stimulus brain activity significantly influences episodic memory formation (Addante et al., 2011; Salari & Rose, 2016; Schneider & Rose, 2016; Scholz et al., 2017; Sweeney-Reed et al., 2016; Winterling et al., 2019). Oscillatory activity, particularly within the theta (3–7 Hz) and alpha (8–12 Hz) bands, has been frequently associated with successful encoding processes (Cruzat et al., 2021; Ostrowski & Rose, 2024). Elevated pre-stimulus theta power has been linked to enhanced binding of contextual information and more accurate source memory, suggesting a preparatory role for subsequent memory performance (Addante et al., 2011). Similarly, increases in alpha oscillations prior to stimulus onset are thought to reflect a state of attentional preparation, facilitating the encoding of complex audiovisual associations (Strunk & Duarte, 2019). Importantly, attentional engagement modulates these oscillatory patterns, indicating that intentional focus can create neural conditions beneficial for memory formation (Schneider & Rose, 2016; Uncapher et al., 2011).

Our findings build upon this established framework, demonstrating that externally induced pre-stimulus alpha entrainment boosts audiovisual associative encoding. The observed increase in alpha power in our experimental condition aligns with theories that posit alpha oscillations as essential for sensory anticipation and attentional gating (Foxe & Snyder, 2011; Kizuk & Mathewson, 2017; Leske et al., 2025; Morrow et al., 2023). This externally driven alpha synchronization likely enhanced preparatory attentional states, enabling more efficient inhibition of irrelevant information and promoting engagement of memory-related neural networks such as parietal and hippocampal regions (Klimesch et al., 2011; Palva et al., 2010; Parish et al., 2018; Raud et al., 2023; Tian et al., 2021). Behaviorally, this facilitation translated into higher recognition sensitivity, driven primarily by increased hit rates, while false alarm rates remained unaffected. These findings support the hypothesis that alpha oscillations modulate sensory preparation and attentional gating during encoding via bottom-up processes, and thereby improve associative memory performance.

In contrast, pre-stimulus theta entrainment did not produce significant behavioral benefits, despite successfully increasing theta power. This suggests that power enhancement alone may be not sufficient to influence memory performance within this paradigm. A key factor could be the temporal specificity of theta’s role in encoding. While theta oscillations were shown to be critical for episodic memory and associative binding (Buzsáki & Moser, 2013; Herweg et al., 2020; Terada et al., 2017), their effectiveness appears to depend heavily on activity during stimulus processing (Hsieh & Ranganath, 2014; Nyhus & Curran, 2010). Prior studies demonstrating memory improvements with theta entrainment typically targeted the period during stimulus presentation, likely optimizing engagement of memory networks (Hanslmayr et al., 2019; Herweg et al., 2020; Köster et al., 2019). These findings indicate that the contribution of theta activity to memory encoding may be more dynamic, occurring during active processing rather than as a preparatory state alone. Our targeting of pre-stimulus activity likely aimed to set a preparatory neural state that may not have directly engaged the neural mechanisms necessary for effective multisensory binding, although oscillatory power was modulated. Furthermore, the role of theta oscillations in memory encoding often involves activity across widespread and synchronized networks such as hippocampal-cortical circuits (Boran et al., 2019; Etter et al., 2023; Nyhus & Curran, 2010), which may not have been fully engaged through unimodal occipital stimulation alone. While the stimulation successfully increased theta power, the lack of phase coherence or cross-regional synchronization may have limited its influence on encoding. These findings emphasize that the contribution of theta activity to memory may be more context-dependent and particularly crucial during active processing phases, rather than solely during pre-stimulus intervals. This aligns with prior studies emphasizing the importance of timing and phase alignment in theta-mediated memory processes.

In addition to the effects of entrainment on primary performance measures we observed a secondary effect on response times. Specifically, participants that received visual stimulation before the encoding of audiovisual pairs responded consistently faster during encoding and, most importantly, during the subsequent recognition task as compared to participants who did not undergo entrainment, while no differences were observed between the three entrainment groups. Given this pattern of results, it is plausible to assume that the faster response times from the entrainment groups could be interpreted as an effect of increased alertness during the encoding period due to general visual stimulation before stimulus onset. Visual stimulation in general has been associated with an increase of alertness before (Figueiro et al., 2018; Golmohammadi et al., 2021; Lok et al., 2018), while alertness, in turn, has been shown to decrease RTs in tasks recruiting executive control systems (Nieuwenhuis & de Kleijn, 2013; Weinbach & Henik, 2012). One could argue that visual information processing might benefit from an improved inhibition of peripheral information (Poirel et al., 2014). This indicates that visual stimulation might have enhanced a preparatory mechanism that is independent from the specific cognitive demand of encoding information but might rather point towards an increased ability to remain vigilant and maintain attention throughout the task despite increasing subjective feelings of fatigue.

Although the entrainment procedure applied in this study led to a reliable modification of pre-stimulus theta and alpha activity, our analyses revealed that both the theta and alpha group exhibited significantly lower beta band power as compared to controls. While we cannot rule out completely that the behavioral effects presented here could also be attributed to modifications of beta band oscillations, it’s plausible to assume that the observed difference was caused by increases in beta activity in the control group rather than decreases in the theta and alpha groups. This is supported by the fact that we found no negative clusters in the beta band when comparing activity from the theta and alpha groups with the NE group, and that the effects common to both the control and NE group contrasts are centered around the respective entrainment frequencies. Instead, the arhythmic stimulation in the control group might have modified pre-stimulus beta-oscillations due to potential additive effects of single-frequency cycles randomly chained together. As individual arhythmic luminance functions were computed for every participant in the control group, individual cycles of the same frequency that ended up at the same time point could have had an amplifying effect during averaging procedures, resulting in what seemed as beta power enhancement. However, this does not invalidate the usefulness of arythmic stimulation, as it plays a complementary role in validating and specifying the precision of entrainment procedures. With this, the present work is in line with previous studies using arhythmic stimulation as an additional control mechanism to ensure that oscillatory responses to the entrainment actually arise from the rhythmicity of a specific frequency (Albouy et al., 2017; Michael et al., 2022; Notbohm & Herrmann, 2016; Thut et al., 2011).

In sum, our results highlight the distinct functional roles of alpha and theta oscillations in multisensory learning and memory. Alpha oscillations appear to serve as a gating mechanism that can be externally modulated to optimize sensory processing and attentional filtering (Foxe & Snyder, 2011; Waldhauser et al., 2012), with our findings providing causal evidence that externally driven alpha rhythms prior to encoding facilitate associative memory performance. In contrast, the unsuccessful behavioral impact of theta entrainment highlights the importance of time specificity and multisensory synchronization for the mnemonic functions of theta oscillations (Herweg et al., 2020; Wang et al., 2018). These insights contribute to a nuanced understanding of how tailored oscillatory modulation can differentially influence neural states underpinning successful memory formation, emphasizing the potential of targeted neurostimulation techniques, personalized cognitive interventions, and novel therapeutic approaches for memory disorders. This is highlighting the significant clinical potential of utilizing specific oscillatory pathways to enhance learning and memory.

## Methods

### Participants

In total, 176 healthy young adults were recruited for this pre-registered study (http://osf.io/5gprt). Participants were required to have normal or corrected-to-normal vision and hearing ability. We had to exclude several participants from the analysis due to unsuccessful entrainment (21.59%). Further exclusions were the result of false positive rates above the predetermined threshold (14.77%). An additional 3.98% were excluded for both failure to entrain and high false positive rate. Taken together, a sample of *n* = 105 (72.38% female) participant data sets were submitted to the analysis, with a group size of *k* = 35 for each experimental group. On average, participants were 24.8 years old (SD = 4.17), with the age ranging from 18 to 35 years. All participants gave their informed consent and received either financial reimbursement or course credit for taking part in the study, which was approved by the ethics committee of the Hamburg Medical Council (PV5893). We confirm that all experiments were performed in accordance with relevant guidelines and regulations.

### Experimental design

The Subsequent Memory Effects task (SME) implemented in this study is a slight variation from the design used in Ostrowski & Rose (2024). The pre-stimulus interval in the encoding task was modified to accommodate the entrainment procedure, while the recognition task remained the same. Participants received the same instructions as in the previous study, with the addition that they were made aware of the presence of an oscillating image. For this study, a between-subjects design was employed, with *entrainment condition* serving as the independent variable with three groups: 5 Hz (theta group), 9 Hz (alpha group), and arhythmic (control group). To take advantage of the Bayesian framework, we implemented an open-ended sequential design for gathering evidence, but added the additional constraint of a maximum group size *k* (Schönbrodt & Wagenmakers, 2018). Thus, data collection was carried out evenly between the groups until a group size with *k* = 15 usable data sets was reached. Subsequent statistical hypothesis testing was conducted incrementally for each additional usable data set, using the pre-registered dependent variable (sensitivity index), with changes in evidence being continuously monitored across all groups. Data collection would stop either when statistical testing showed moderate support for either the alternative or null hypothesis (*BF*_10_ > 3 or *BF*_10_ < 1/3; Jeffreys, Harold, 1998; Lee & Wagenmakers, 2014) or when group size reached *k* = 35 for each group. This resulted in group sizes of *k* = 35 for the theta, alpha, as well as the control group.

### Stimulus material

Stimulus pairs consisting of one image and one sound were selected randomly from an internal database, and the selection was unique for each experimental run. All images featured a resolution of 640 x 480 pixels and a 24-bit color depth. Each image depicted a photograph of either natural or man-made scenes. An additional neutral image depicting a photograph of Earth in space was chosen as the entrainment stimulus to be shown in every trial. We inverted the colors of the entrainment stimulus to increase contrast, thereby increasing the intensity of the stimulation. According to the principle of the Arnold tongue (Pikovsky et al., 2003; Tass et al., 1998), higher stimulation intensity might compensate for a slight frequency mismatch between the entraining signal and the ongoing oscillations in the brain, thus increasing the probability of a successful entrainment. The sounds were real-life recordings of either sounds from nature (e.g. animal calls) or from man-made or artificial environments (e.g. a honk of a car). All sounds were cropped to a duration of 2 s, and featured a bit rate of 1411 kBit/s. All pairings were created in a manner so that no effects of semantic congruency would arise (Parise & Spence, 2012). While it was possible that e.g. animal images could be paired with animal sounds, pairings containing an image of an animal and the corresponding sound of that animal were excluded.

### Sensory entrainment

Sensory stimulation was used during the pre-stimulus intervals of encoding trials to manipulate narrow-band oscillatory activity and investigate its effects on subsequent encoding. The entrainment stimulus, which was the same for all participants and all groups, was presented for 2 s before stimulus onset. Specifically, its luminance varied in a sinusoidal manner, resulting in a rhythmic oscillation of 5 Hz in the theta group and 9 Hz in the alpha group. To achieve high temporal resolution of the luminance sine curve, we used a monitor with a frame rate of 240 Hz (Alienware 27 AW2723DF, Dell Technologies, Round Rock, USA). This enabled us to change luminance every 4.2 ms, resulting in luminance change that closely followed a sine curve instead of a box car function. The frequencies of the entrainment signal were determined based on evidence from our previous study (Ostrowski & Rose, 2024), where the peak subsequent memory effects in the pre-stimulus interval were found at 5 Hz in the theta range, as well as 9 Hz in the alpha range. While the luminance in the theta and alpha groups was kept at a steady rhythm in the respective frequencies, the luminance waveforms in the control condition were arhythmic. The waveforms consisted of single cycles of differing frequencies pulled randomly from the interval of 13 to 24 Hz. Importantly, we excluded frequencies of 15 Hz, 18 Hz, and 20 Hz, as these are harmonic frequencies of the entrainment frequencies in the other entrainment conditions. Each participant in the control group was presented with a unique arhythmic waveform with a duration comparable to the 2 s of entrainment in the other groups (M = 1.976 s, SD = 0.022). Notably, the luminance waveform for every group always started and ended at zero luminance (image not visible). To ensure that the stimulus pair would be presented in line with the entrainment rhythm, we implemented a gap of 100 ms between the end of stimulation and stimulus onset in the theta group (56 ms in the alpha group, respectively), which constitutes half of a cycle in the entrainment frquency. In the control condition, this gap was set to 50 ms.

### Task and procedure

The experimental procedure was the same regardless of experimental group. After giving informed consent and receiving a short introduction by the experimenter, participants were seated in a sound-attenuated chamber. The experimental session started with a recording of 3.5 minutes of resting-state activity, during which participants were told to fixate a fixation cross on the screen. This was followed by the SME task, which consisted of a short training session and three experimental runs that only differed in the stimulation material presented to the participants. Each experimental run included an encoding phase, an intermission, and a subsequent recognition phase (see **Figure 1**). One encoding phase consisted of 47 trials. During each trial, participants were simultaneously presented with an image and a sound for 2 s. A white fixation cross was visible during stimulus presentation and remained on the screen for 3 to 5 s after stimulus offset. Before stimulus onset, the entrainment stimulus was presented in the respective frequency. Participants were instructed to memorize the combination of image and sound from every trial. Furthermore, participants should indicate whether both the image and sound represented an animal (right mouse button) or not (left mouse button). Button presses were registered as a valid response during the first 3 s after stimulus onset but were otherwise counted as a missed response. The experimental trials were followed by four survey questions measuring the participants’ perception of the entrainment procedure. Specifically, the questions measured salience, attention, fatigue, and distractive qualities in relation to the entrainment procedure. During the subsequent intermission of approximately 3 minutes, the participants were asked to count down aloud from 100 (115 and 125 in the second and third run, respectively) in steps of 7 (9 and 13 in the other runs, respectively).

In the recognition phase, the 47 audiovisual pairings from the preceding encoding phase were presented again but intermixed with 47 new pairings, which were created by randomly shuffling the original ones. Note that the individual images and sounds used for the combinations remained the same within each experimental run. All stimulus pairs were again presented for 2 s, with a small white fixation cross layered on top of the image. The fixation cross remained on the screen after stimulus onset. The participants were asked to indicate via button-press whether the current pair had already been presented in the preceding encoding phase (left mouse button) or not (right mouse button). They were further encouraged to press the right mouse button when they felt highly uncertain about a stimulus pair. As in the encoding phase, valid responses were recorded up to 3 s after stimulus onset, and otherwise labeled as a missed response trial. The subsequent inter-trial interval was set to 4 s, during which the white fixation cross was visible on the screen. Across all three experimental runs, participants were presented with 141 unique encoding trials and 282 recognition trials. At the end of the experimental session, resting-state activity was measured again for 3.5 minutes while participants fixated the middle of the screen.

### EEG data acquisition and preprocessing

We used a 64-channel electrode setup (ActiCap, BrainProducts, Gilching, Germany) to record EEG. Four of those electrodes were placed on the left and right temple, as well as above and below the left eye, to record vertical and horizontal EOG. The signal was referenced online to *FCz* and re-referenced offline to a common average. The ground electrode was placed at *Iz* below *Oz*, and electrode impedences were kept below 10 kΩ. The signal was amplified with a low cut-off frequency of 0.53 Hz (0.3 s time constant) and recorded at a sampling rate of 500 Hz. EEG activity was recorded during all encoding and recognition phases, but not during intermissions. These settings were used for resting-state recordings as well as for the recordings during the SME task.

Offline preprocessing was done using the Fieldtrip (Oostenveld et al., 2011) and EEGLAB (Delorme & Makeig, 2004) toolboxes for MATLAB (Release 2023a, The Mathworks Inc., Natick, Massachusetts, USA). For the data from the encoding task, an automated approach was used to epoch and clean the data for further processing. The raw data was divided into segments from –3.4 s to 2.5 s relative to the onset of the stimulus pair. A bandpass filter was used to filter out all frequencies outside the range of 0.5 Hz to 40 Hz. Next, trials containing temporally distinct artifacts based on muscular activity or related to electronics were rejected in an automated pipeline using the *ft_artifact_zvalue* function from Fieldtrip. The trial data was filtered, z-transformed, and averaged over channels. An accumulated z-score was computed for each trial based on the types of artifacts. The cutoff value was set to z = 60 for jump artifacts and z = 30 for artifacts caused by phasic muscular activity. Trials were then rejected if the accumulated z-score was larger than the corresponding threshold value. The resulting data was submitted to an automated Independent Component Analysis (ICA) to remove underlying noise from muscular activity as well as artifacts resulting from blinks and eye movements using the *ICLabel* plugin for EEGLAB (Pion-Tonachini et al., 2019). Components that showed at least a probability of 80% of being related to eye-movements, noise caused by muscular activation, or line noise were flagged for removal. On average, 6.37 (SD = 3.8) independent components were removed from the data of the theta group, 6.17 (SD = 3.99) for the alpha group, and 6.63 (SD = 3.32) for the control group. The data was then re-referenced again to the common average. After preprocessing, 4.74 trials (SD = 4.9) out of 141 encoding trials were removed from data sets in the theta group. In the alpha group, an average of 5.14 trials (SD = 3.84) was rejected per participant, while 3.77 trials (SD = 2.65) were rejected in the control group. As the data from the previous study were also analyzed again in the context of the present investigation (Ostrowski & Rose, 2024), all corresponding EEG data were submitted to the same processing pipeline to ensure comparability. After ICA, 4.79 (SD = 2.19) independent components were rejected from the data on average per participant. After preprocessing, an average of 6.42 trials (SD = 4.67) per participant was removed from the data.

From the 3.5 minutes of recorded resting-state activity before and after the experiment, the first and last 15 seconds were omitted for offline processing. Pre– and post-experiment data were processed separately. From this point, we will refer to the data as RestPre and RestPost, respectively. We used a bandpass filter to remove activity below 0.5 Hz and above 40 Hz from the remaining 3-minute interval. The data were then divided into 90 epochs with a length of 2 s each and cleaned from temporally distinct artifacts with the same automated pipeline that was used with the experimental data. Epochs containing artifacts were then removed from the data. The results were submitted to an automated ICA using the same parameters as for the experimental data. On average, 1.11 epochs (SD = 1.3) were removed from RestPre data in the theta group per participant (alpha: 0.91, SD = 1.07; control: 1.2, SD = 1.45). After ICA, 2.91 (SD = 1.8) independent components were rejected per participant in the theta group (alpha: 2.97, SD = 2.35; control: 3.17, SD = 2.16). For RestPost, an average of 1.4 epochs (SD = 2.24) were removed in the theta group per participant (alpha: 0.89, SD = 1.47; control: 1.14, SD = 1.57). On average, 3.74 (SD = 2.78) independent components were rejected (alpha: 3.54, SD = 2.76; control: 5.51, SD = 3.78).

### Entrainment validation

As a first step, the pre-processed experimental data from the theta and alpha groups were decomposed into the time-frequency domain. We chose a frequency range of 1 to 40 Hz with frequency bins of 1 Hz, and a time interval of –3.1 s to 2.2 s relative to the onset of the stimulus pairs. Fieldtrip’s *mtmconvol* method (Oostenveld et al., 2011) was used in conjunction with a Hanning window of 500 ms and a step size of 100 ms. The additional 300 ms before and after the chosen time interval that were retained during preprocessing served as padding to avoid edge artifacts from the decomposition process. After conducting the decomposition for every trial, the resulting oscillatory power was then averaged over trials for every participant. Next, the data was normalized using a measure of change percentage relative to baseline activity that was defined as the activity from –3.1 s to –2.1 s before stimulus onset. For every individual data set from the theta and alpha group, an average was computed from the data of occipital electrodes (O1, O2, and Oz). As the sensory entrainment took place in the visual domain, the most prominent response should be expected in the electrodes adjacent to the visual cortex. Separate frequency envelopes were chosen for the theta group (5 Hz ± 1 Hz) and the alpha group (9 ± 1 Hz), with a common time interval of interest ranging from –1.1 s to – 0.1 s relative to stimulus onset. We used the latter half of the entrainment interval to estimate entrainment success, as phase alignment and entrainment typically develop over time and tend to plateau after an initial adjustment period (Riecke et al., 2015; Wacker et al., 2011). Assessing the full interval may underestimate entrainment strength due to lower power at the beginning of the stimulation. The entrainment was deemed successful if a relative change in power of at least 10 % could be observed for at least 500 ms within the time interval of interest.

## Statistical analysis

### Behavioral data

We used a Bayesian framework to test the hypotheses relating to behavioral effects, utilizing the *BayesFactor* package for *R* (v.4.3.3). In line with the signal detection theory (Pastore & Scheirer, 1974; Stanislaw & Todorov, 1999), four percentage measures were extracted for every participant from the recognition data: Correctly remembered old pairings (*hits*), not remembered old pairings (*misses*), new pairings correctly rejected as new (*correct rejections*), and new pairings seemingly remembered as old (*false alarms*). Our main dependent variable, the sensitivity index *d’*, was computed by calculating the difference between the z-transformed hit and false alarm rates for every participant. When group size reached *k* = 15, we used the Bayesian version of a *t*-test to statistically compare memory performance between groups. Testing was then repeated every time *k* increased by one for each group. Specifically, one-sided tests were computed to compare performance between the theta group and controls (H2a), as well as between the alpha group and controls (H2b). For estimating differences between both entrainment groups (H2c), a two-sided test was performed. In all cases, a Cauchy distribution of medium width was used as prior, i.e. with an *r* scale of √2/2.

For the comparison of the experimental groups from the current study with the data from the previous investigation (H3), the NE group data was processed in the same manner. To keep in line with our exclusion criteria, participants with a false positive rate > 25% were not considered in the analysis, resulting in a sample size of k = 45 for the NE group. Due to the difference in group size, a sampling approach was chosen in which a subsample was randomly pulled from the NE data set that matched the group size of the entrainment groups. The average sensitivity was calculated from that subsample and compared to the mean of the full NE sample. This procedure was repeated 50 times. Ultimately, we chose the subsample where the difference in means was minimal, ensuring that the subsample would be representative of the original NE sample. We then conducted one-sided Bayesian t-tests to compare memory performance between both entrainment groups and the previous data set using the same settings as in testing for H2.

Bayesian statistical approaches were further used to explore differences between groups in secondary behavioral variables. A Bayesian one-way ANOVA with the factor *entrainment condition* (theta, alpha, control, NE) was used to assess differences in accuracy and response times from the categorization task during encoding. Changes in memory sensitivity over the course of the experiment were investigated using a mixed-design Bayesian ANOVA with the factors *entrainment condition* and *experimental run* (a, b, c). Furthermore, Bayesian one-way ANOVAs with the factor *entrainment condition* were used to assess group differences for every response category of reaction times. To investigate, whether the entrainment condition modulated the discrepancy in response time between correct and incorrect trials, a mixed-design Bayesian ANOVA with the factors *entrainment condition* and *correctness* was utilized. Finally, Bayesian mixed-design ANOVAs with the factors *entrainment condition* and *experimental run* were used to assess differences in the subjective perception of the entrainment procedure, as well as state of attention and fatigue. The analyses were conducted separately for each survey item. To estimate the relative likelihood of the interactions in the these analyses, the ratio of Bayes factors corresponding to the full model and the model containing only the main effects was computed. For all analyses, a Cauchy distribution with an *r* scale of √2/2 was used as prior.

### EEG data

To statistically test the success of entrainment, we compared oscillatory activity from the theta and alpha groups with activity from the control group. Specifically, the baseline-normalized time-frequency data was restricted to the latter half of the entrainment period (−1.1 s to –0.1 s relative to stimulus onset), and the frequency range was set to 1 to 40 Hz. We used a non-parametric permutation testing approach with a cluster-based correction for multiple comparisons as implemented in Fieldtrip (Oostenveld et al., 2011). Independent-samples *t-*tests were computed for every data point across participants from the channel-time-frequency space. Data points that showed significant differences between conditions (*p* < .05) were organized into clusters based on temporal, spatial, and spectral proximity. For each cluster, statistical values were summed to yield a cluster-level statistic, and the highest of these sums was selected as the principal test statistic for condition comparisons. To construct a reference distribution, a Monte Carlo approach was employed: all trials from both conditions were merged into a single dataset and randomly split into two groups. Statistical testing was performed again at the level of individual data points within these shuffled groups, and cluster-level statistics were recalculated. This randomization process was repeated 4000 times. During each iteration, the largest cluster-level statistics were recorded to generate the null distribution, separately for positive and negative clusters. The final *p*-value for condition differences was obtained by determining the proportion of randomizations that produced a test statistic greater than that observed in the original data. This method was applied across all detected clusters, yielding a *p*-value for each cluster’s comparison between conditions. The same statistical approach was used for the comparison of pre-stimulus activity from the theta and alpha groups to activity from the NE group. In addition, this approach was also used to assess differences between the entrainment groups (theta, alpha, control) in brain activity during stimulus presentation. However, an independent-samples *F*-test was used on sample level in this case.

A similar statistical approach was used for the exploration of resting-state data. For every entrainment group, the preprocessed RestPre and RestPost data were decomposed into the frequency domain by using the Fast Fourier Transform on single epochs for a frequency range of 1 to 40 Hz. All epoch spectra were then averaged to a subject-specific mean frequency spectrum. This was done separately for RestPre and RestPost. For each of the entrainment groups, we compared RestPre and RestPost with a paired-samples *t-*test on sample level. To further assess whether these differences varied between groups, difference scores were computed between RestPre and RestPost data and submitted to cluster-based permutation testing using an independent-samples *F*-test on sample level. Thresholds for *p*-values were kept as laid out above.

To determine each participant’s Individual Alpha Frequency (IAF) and Individual Theta Frequency (ITF), we analyzed resting state EEG data acquired prior to the experimental task. Preprocessed data underwent spectral analysis using a multitaper Fast Fourier Transform (FFT) approach implemented in FieldTrip. Spectral power was computed across frequencies from 1 to 40 Hz in 1 Hz steps, with a 2 Hz smoothing kernel applied. For each participant, we extracted the mean power spectrum across a set of posterior electrodes (Pz, POz, Oz, O1, O2, P3, P4, PO3, PO4). The IAF was identified as the frequency within the alpha range (8–12 Hz) exhibiting the maximum spectral power across these electrodes. Conversely, the ITF was defined as the frequency within the theta range (3–7 Hz) with the highest spectral power. We calculated the absolute differences between each participant’s IAF and ITF and their corresponding entrainment frequencies (theta: 5 Hz; alpha: 9 Hz). To assess whether the proximity of an individual’s intrinsic frequencies to the stimulation frequencies influenced entrainment efficacy, we conducted Pearson correlation analyses between these frequency-distance measures and the maximum relative change in spectral power at the individual peak channel during stimulation.

## Data availability

The raw EEG and behavioral data underlying our findings have been uploaded to an open repository of the University of Hamburg for accessibility (https://doi.org/10.25592/uhhfdm.17620).

## Acknowledgements

The present research was supported by the German Research Foundation (DFG; SFB/Transregio 169, Project B3 and DFG RO 2653/9-1). We thank Michele Frerichs and Lennart May for assistance during data acquisition.

## Author Contributions

M.R., J.O. and M.M. designed the study. J.O. and M.M. performed data acquisition. J.O. and M.M. analyzed the data. M.R. acquired funding, conceptualized, and supervised the project. J.O., M.M. and M.R. wrote the original manuscript. J.O., M.M., and M.R. reviewed and edited the manuscript and approved the final manuscript.

## Additional Information

Correspondence and requests for materials should be addressed to M.R.

## Supplementary material

**Supplementary 1. Contrasting pre-stimulus activity from theta and alpha groups to activity from the control and NE groups**

**Figure S1.**
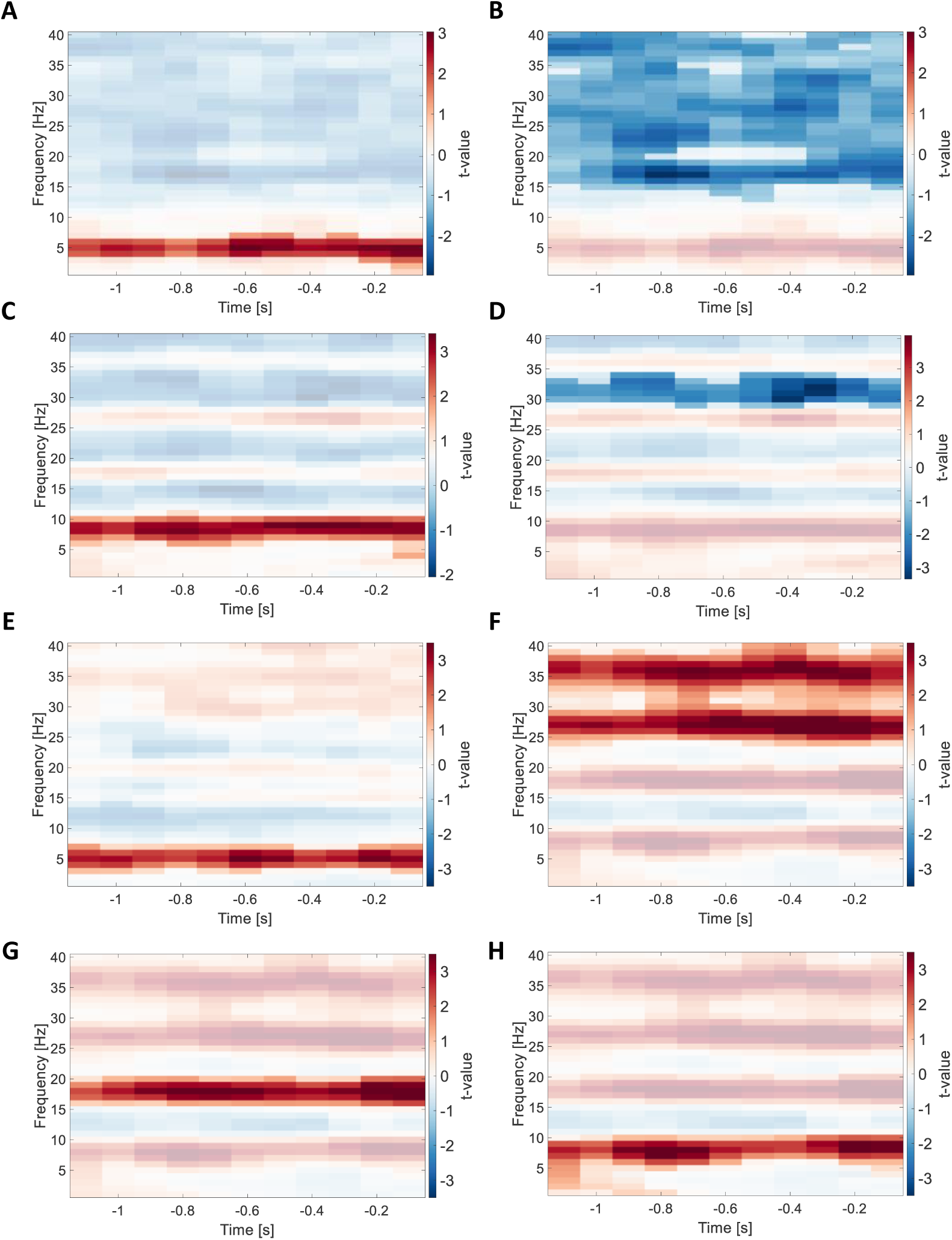
Results of EEG data contrasts of entrainment groups with the control and NE groups. The figure shows time-frequency plots depicting the results of the statistical comparison of relative change in pre-stimulus activity. **(A)** and **(B)** show the two significant clusters from the comparison of the theta group with the control group. **(C)** and **(D)** depict the two significant clusters resulting from contrasting the alpha group with the control group. **(E)** shows the positive cluster revealed by comparing activity from the theta group with the NE group. **(F) – (G)** depict the statistical results comparing the alpha group with the NE group. In all time-frequency plots, positive *t*-values signify greater relative change in the theta or alpha groups, respectively. Opaque data points show the extent of a statistically significant cluster (*p* < .025, corrected). Each subplot shows one distinct cluster and depicts the *t*-values averaged over the electrodes comprising the cluster.

Oscillatory power in the late entrainment period (−1.1 s to –0.1 s relative to stimulus onset) from the theta and alpha group was each contrasted with the activity from the NE group in the same time period. We used a cluster-based permutation approach to account for multiple comparisons, and two-tailed independent-samples *t*-tests on sample level. The frequency range for this analysis was set to 1 to 40 Hz, and all electrodes were included. Comparing the theta entrainment condition with the NE group, the analysis yielded one significant positive cluster in the frequency range of 3 to 7 Hz, spanning the whole analysis window (*p* < .025, corrected). This suggests significantly increased oscillatory power in the envelope around 5 Hz for the theta group as compared to the NE group **(Figure S2E)**. Contrasting activity from the alpha group with the NE group revealed three distinct positive clusters, each spanning the whole analysis window. The clusters covered the frequency ranges of 24 to 40 Hz and 16 to 20 Hz. Importantly, the third cluster ranged from 1 to 10 Hz up until –0.8 s relative to stimulus onset and was centered one the 9 Hz envelope for the remaining part of the analysis time window (**Figure S2H**).

**Supplementary 2. Analysis of oscillatory activity in the stimulus presentation window during encoding**

**Figure S2.**
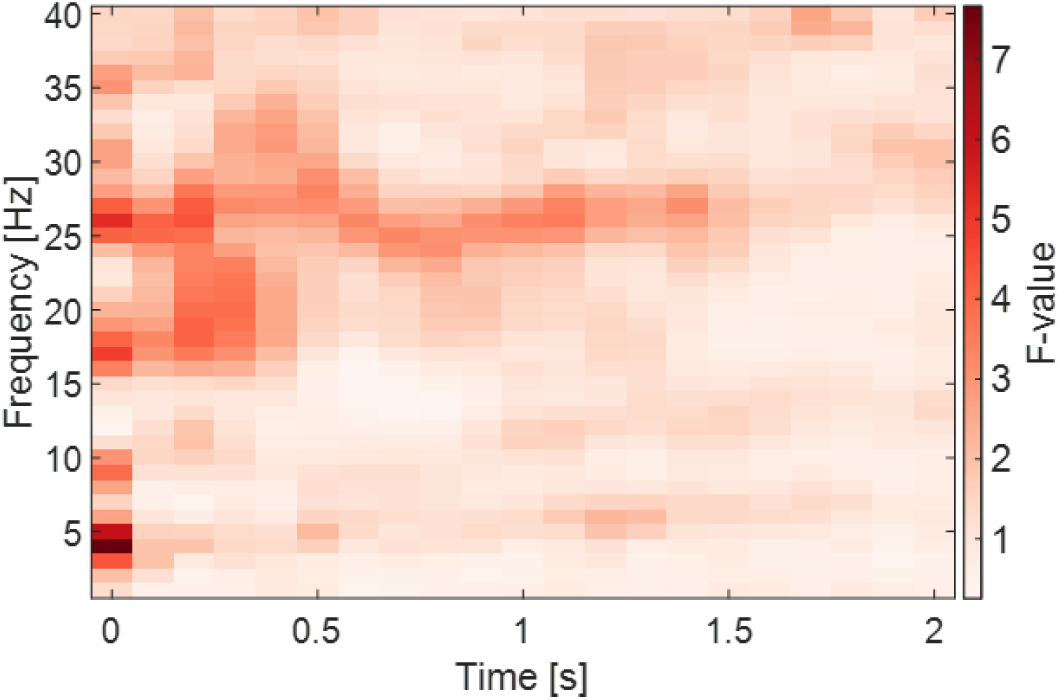
Difference in post-stimulus activity between entrainment groups. The figure shows a time-frequency plot of the stimulus presentation time window from the encoding task over a frequency range of 1 to 40 Hz. The color dimension displays the F-values from the independent-samples F-test.

In order to assess potential differences in oscillatory activity during stimulus presentation between groups, we compared oscillatory power from the post-stimulus interval (0 s to 2s relative to stimulus onset) among the entrainment groups (theta, alpha, and control) using an independent-samples *F*-test on sample level. Data was included for a frequency range of 1 to 40 Hz and all electrodes, and cluster-based permutation was used for multiple-comparison corrections. However, the analysis showed only a tendency for a significant cluster in the electrode-frequency-time space, suggesting that there are no significant differences in post-stimulus power between the entrainment groups (*p* = .069, corrected).

**Supplementary 3. Analysis of categorization task performance during encoding**

**Figure S3.**
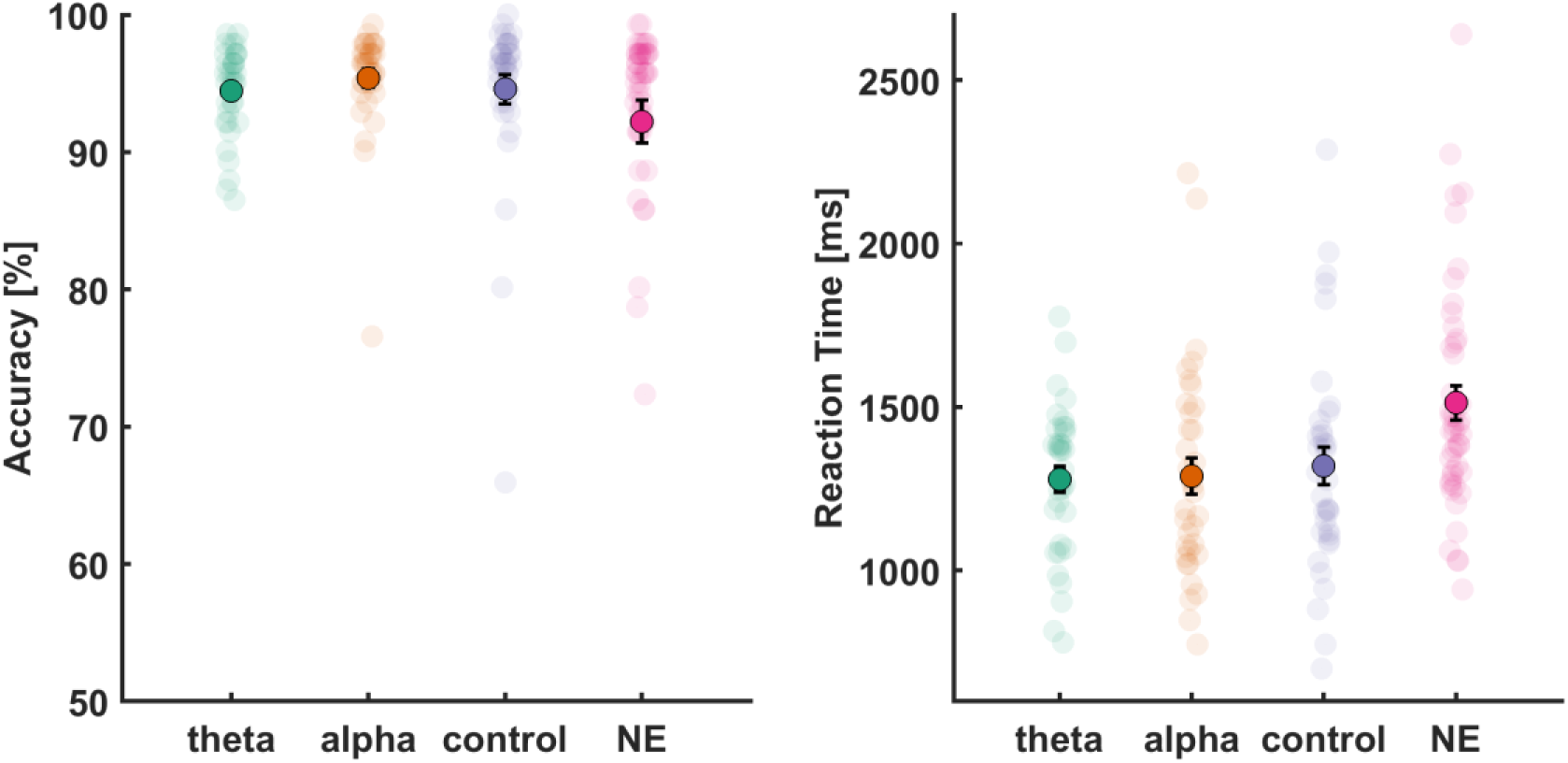
Performance in the categorization task during encoding. *Left*: Accuracy values for the categorization task during encoding for each group. *Right:* Average response time for the categorization task during encoding. Transparent data points mark individual task accuracy, and the black error bars signify the standard error of means.

In the categorization task from the encoding phase, participants showed high accuracy across all entrainment conditions, 94.45% (SD = 3.23) for theta, 95.38% (SD = 3.88) for alpha, and 94.57% (SD = 6.27) for control condition. Accuracy in the NE condition as an additional control was slightly lower at 92.21% (SD = 10.43). The overall accuracy across all four conditions was 94.02% (SD = 6.97). Using a Bayesian one-way ANOVA model yielded a Bayes factor of BF₁₀ = 0.381, suggesting moderate evidence in support of the null hypothesis of no significant differences among the groups. These results suggest that participants in the current study maintained high compliance with task demands throughout the experiment, which was essential for accurately assessing the subsequent impact of oscillatory activity on memory performance. Reaction times showed a similar pattern. Participants responded fastest in the entrainment conditions (theta: 1279.4 ms, SD = 230.3; alpha: 1288.9 ms, SD = 327.6; control: 1320.4 ms, SD = 343.7), with slower responses in the NE condition (1512.6 ms, SD = 355.3). The overall average reaction time was 1361.11 ms (SD = 333.1). However, a Bayesian one-way ANOVA indicated moderate evidence for the alternative hypothesis, BF_10_ = 5.0737, indicating measurable differences among the groups. Individual group contrasts revealed that there was likely no difference in response times between the entrainment groups (theta vs control: BF_10_ = 0.2849; alpha vs control: BF_10_ = 0.2628; theta vs alpha: BF_10_ = 0.248). However, the evidence suggests a moderate-to-strong effect for differences between the theta and alpha groups and the NE group (theta vs NE: BF_10_ = 19.1684; alpha vs NE: BF_10_ = 5.7022). The comparison between the control and the NE group yielded only weak evidence for significant difference, BF_10_ = 2.3366.

**Supplementary 4*. Changes in performance over the course of the experiment***

**Figure S4.**
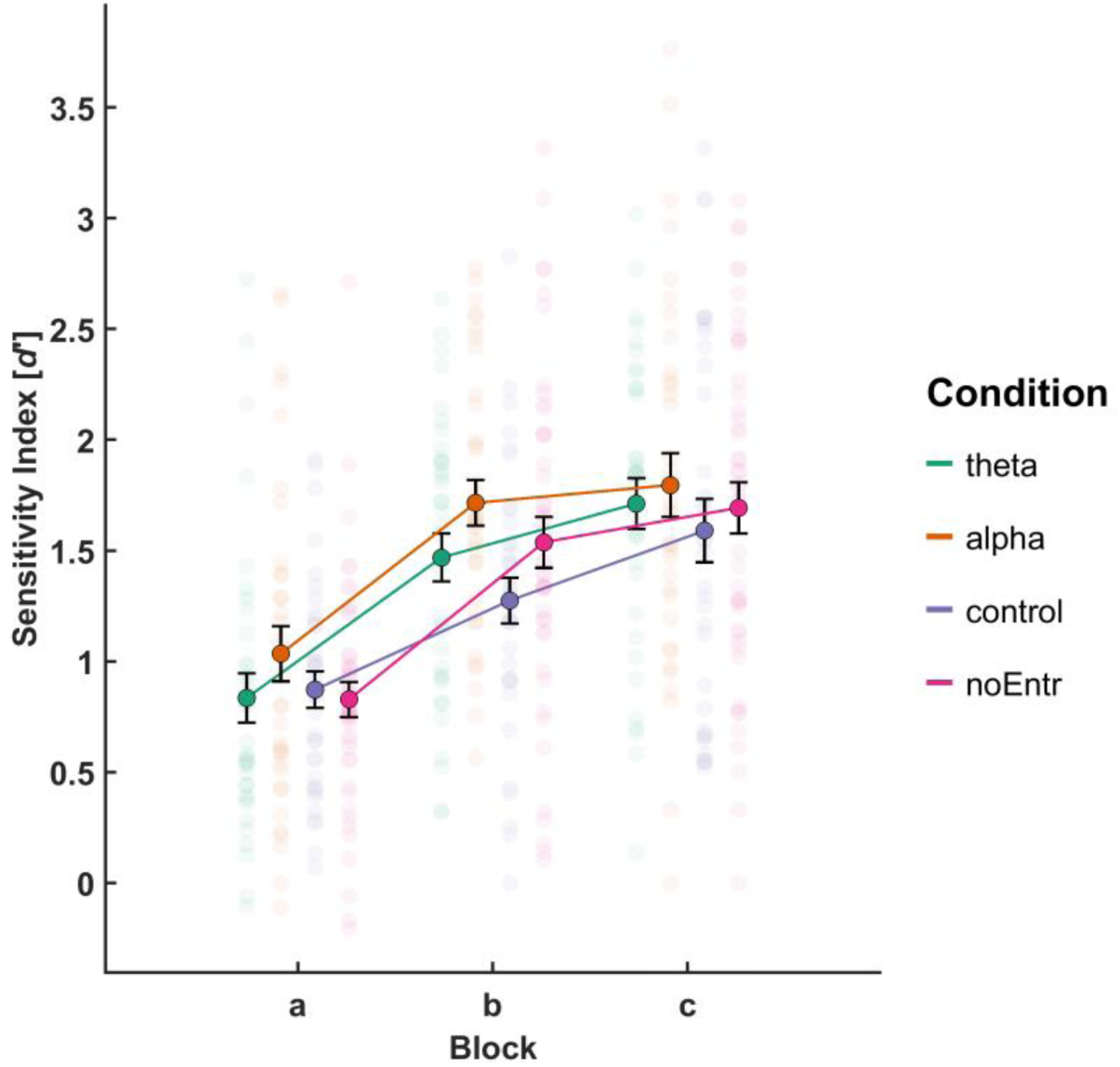
Changes in sensitivity indices across experimental runs. The plot depicts mean sensitivity indices (*d’*) over participants for every group and the three experimental runs. Every transparent data point marks the individual sensitivity index of one participant. Black error bars indicate the standard error of means.

To assess changes in behavioral performance over time, we conducted a mixed-design Bayesian ANOVA with the within-subjects factor *block* (a, b, c) and the between-subjects factor *entrainment condition* (theta, alpha, control, NE), both as fixed factors. The participant ID was included as a random effect. The best-supported model included *block* and the participant ID, BF = 1.58 × 10³⁵, indicating extreme evidence for a main effect of *block*. Adding *entrainment condition* reduced model support by a factor of approximately 4.6 (BF = 3.43 × 10³⁴), while including the *block* x *entrainment condition* interaction further reduced support by a factor of approximately 135, BF = 1.17 × 10³³. The model with only *entrainment condition* and participant ID was 7.3 times less likely than the null model (BF = 0.14), providing strong evidence against a main effect of *entrainment condition*. This indicates that the improvement in memory performance over the course of the experiment was consistent across entrainment conditions and was not modified by the type of entrainment.

***Supplementary 5*. Individual group and variable contrasts for response times during recognition**

**Table S1.**
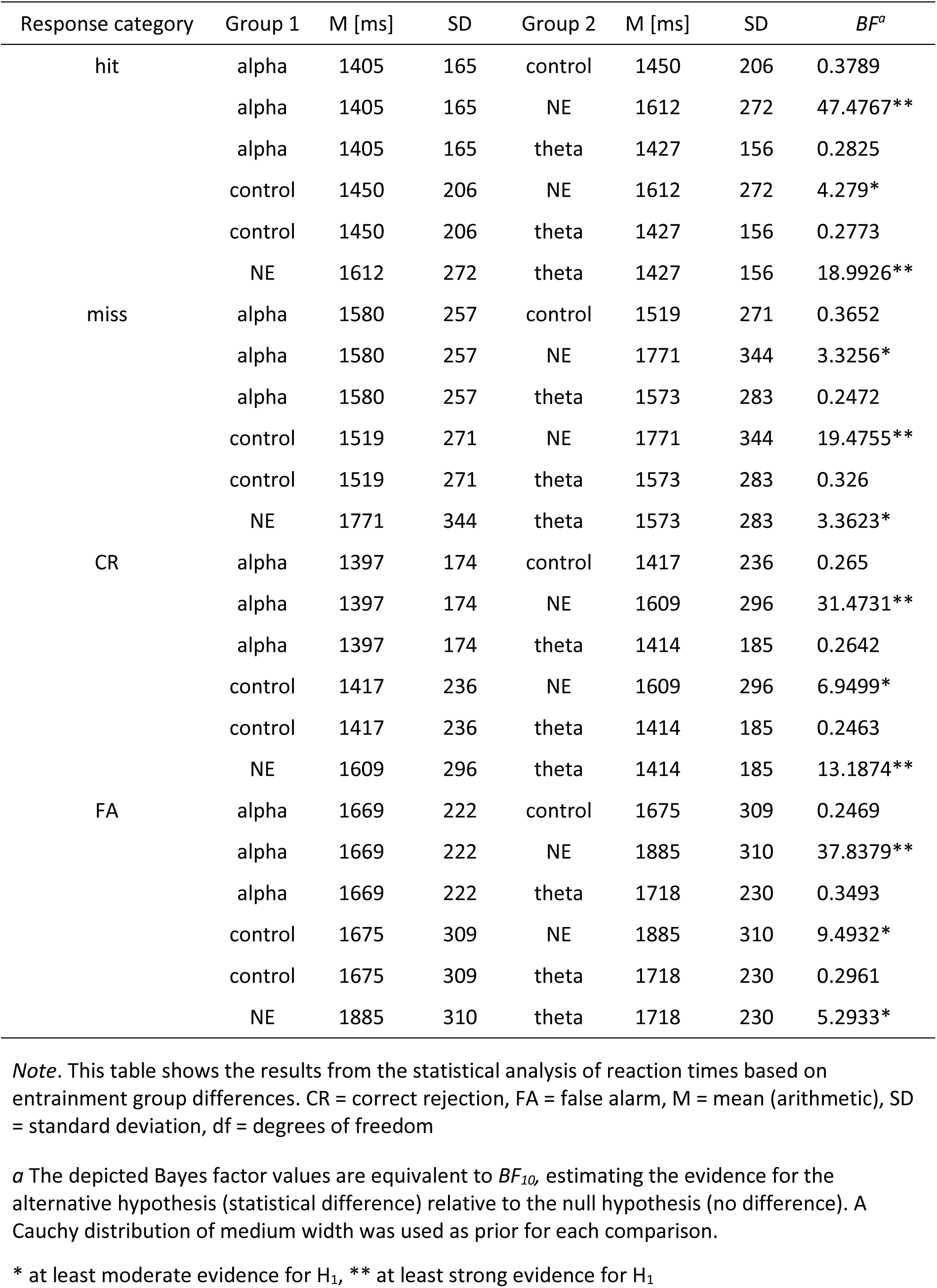
Individual group comparisons of RTs for all response categories.

**Supplementary 6*. Resting State***

**Figure S5.**
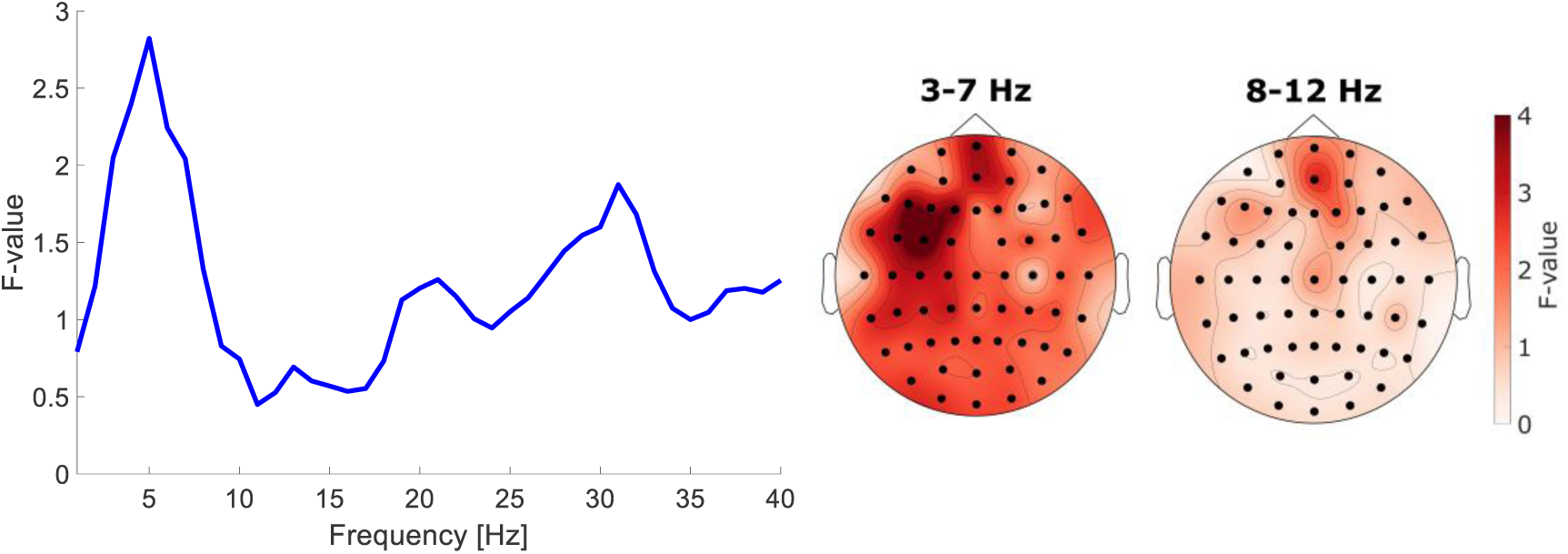
Group contrast for differences in resting-state activity before and after the experiment. *Left*: F-values for the analysis window of 1 to 40 Hz from a cluster-based permutation test with an independent-samples *F-*test on sample level assessing group differences in pre– and post-experiment resting-state discrepancies. *Right*: Topographical distribution of *F*-values averaged across the 3–7 Hz (theta) and 8–12 Hz (alpha) bands. No significant clusters were observed in this analysis (p = .2972, corrected).

As some studies report lingering oscillatory effects due to entrainment procedures **[Kasten & Herrmann, 2022; Gallina et al., 2023)**, we explored differences in resting-state spectra that were recorded once before (RestPre) and once after the SME task (RestPost) to determine whether traces of the entrainment could be observed even after the experiment. For the analysis, we used power spectra in the frequency range of 1 to 40 Hz and submitted the data to a cluster-based permutation test with two-tailed paired-samples t-tests on the sample level. Note that this analysis was conducted separately for the theta group, alpha group, as well as the control group. For the theta group, one negative cluster was found in the alpha band (8 to 12 Hz), indicating increased power after the experiment (p < .025, corrected). Similarly, a negative cluster ranging from 7 to 18 Hz was observed for the comparison in the alpha group (p < .025, corrected), while the analysis in the control revealed a negative cluster in the alpha band (8 – 12 Hz, p < .025, corrected). Interestingly, the control group analysis yielded a second negative cluster in the beta band, ranging from 17 to 33 Hz (p < .025, corrected). As the effect in the alpha band and, to a certain degree, in the beta band was observed in all three groups, we were interested in whether the effect magnitude differed between the groups.

**Supplementary 7*. No differences in subjective perception of entrainment***

Survey items (translated from German into English):

*I1 (pleasentness)*: How pleasant did you find the flickering of the image?

*I2 (distraction)*: To what extent did you feel distracted by the flickering of the image while trying to remember the pairs?

*I3 (attention)*: How would you rate your level of attention during the task?

*I4 (fatigue)*: How exhausted do you feel at the moment?

Participants rated on a scale from 0 (*not at all*) to 5 (*very much*) in steps of 0.5.

**Figure S6.**
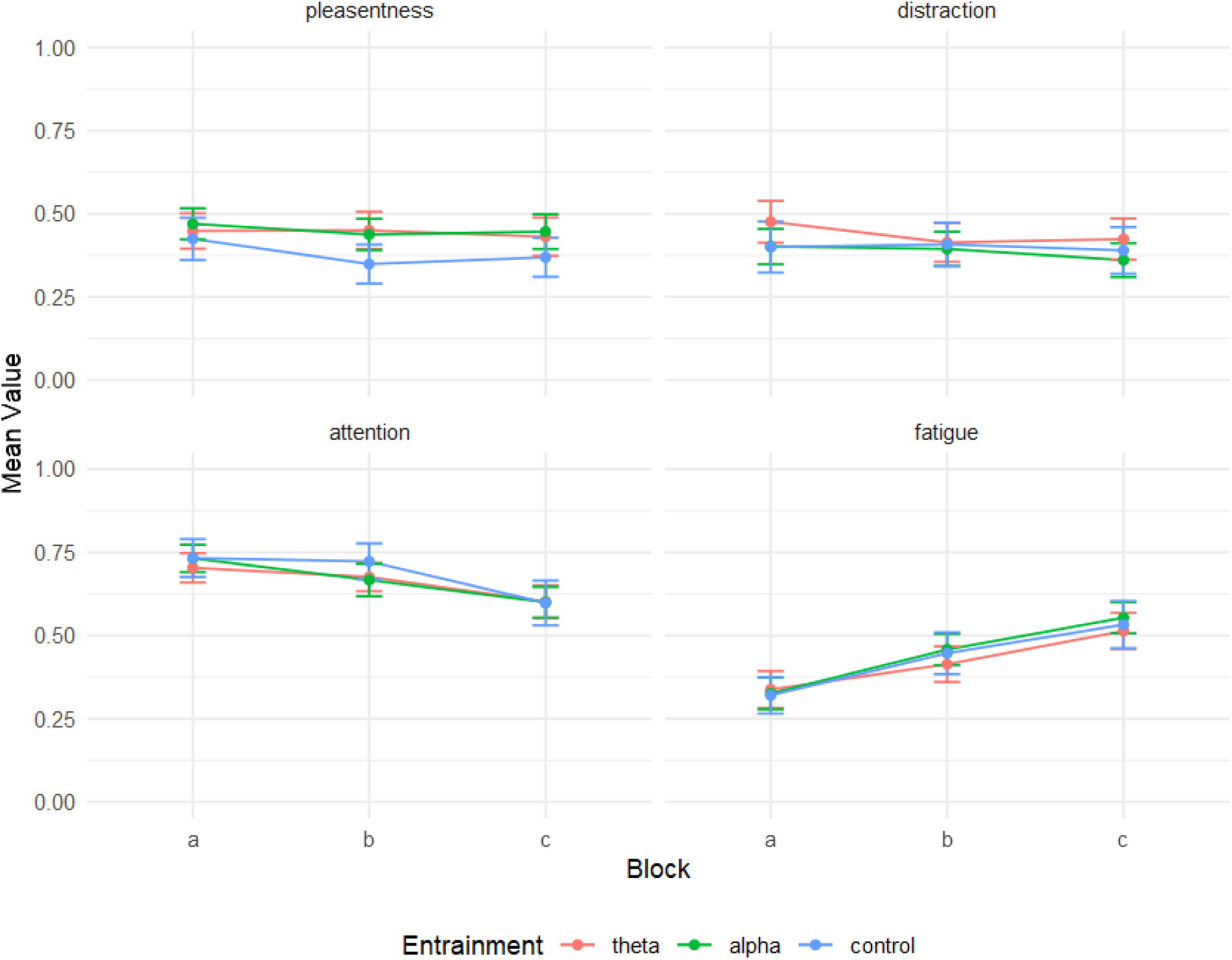
Subjective ratings of task-related experience across blocks and entrainment conditions. The data points depict group and block averages across participants. The error bars mark the standard error of means.

As individual perception qualities of images with oscillating luminance may vary, we investigated whether the subjective perception of the sensory stimulation might differ between the entrainment groups to control for salience effects. Participants received four survey items after each encoding phase, measuring the *pleasantness* and the *distracting qualities* of the entrainment procedure, as well as *attention* and *fatigue*. We conducted a Bayesian mixed-design ANOVA for every item, with a between-subjects factor *pre-stimulus condition* (theta, alpha, control) and a within-subjects factor *block* (A, B, C). Scores from every item did not significantly differ between levels of *pre-stimulus condition* (BF_pleasentness_ = 0.3099, BF_distraction_ = 0.3853, BF_attention_ = 0.1132, BF_fatigue_ = 0.1279), indicating that the type of entrainment procedure had no differential effect. However, the analysis revealed an effect of *block* for the variables *distraction, attention,* and *fatigue* (BF_distraction_ = 7.8834, BF_attention_ = 1.855 x 10^9^, BF_fatigue_ = 1.3415 x 10^15^). No interactions of *pre-stimulus condition* and *block* were observed (BF_pleasentness_ = 0.0732, BF_distraction_ = 0.0578, BF_attention_ = 0.195, BF_fatigue_ = 0.0552. Participants felt less distracted by the entrainment in block C of the experiment (M = 0.382, SD = 0.224) than in block A (M = 0.45, SD = 0.243). Conversely, participants rated their level of attention in block C (M = 0.573, SD = 0.203) consistently lower than in block A (M = 0.694, SD = 0.186). This was accompanied by increased fatigue ratings in block C (M = 0.528, SD = 0.223) as compared to block A (M = 0.338, SD = 0.197). In sum, evidence from the survey data indicates that the entrainment procedures were received equally pleasant and distracting, suggesting no confound of the behavioral results due to subjective perception.

