## Supplemental Table 1 for "Disentangling the Functional Roles of Pre-Stimulus Oscillations in Crossmodal Associative Memory Formation via Sensory Entrainment"

**Table S1**

*Individual group comparisons of RTs for all response categories*

| Response category | Group 1 | M [ms] | SD | Group 2 | M [ms] | SD | *BF*[a] |
|---|---|---|---|---|---|---|---|
| hit | alpha | 1405 | 165 | control | 1450 | 206 | 0.3789 |
| | alpha | 1405 | 165 | NE | 1612 | 272 | 47.4767** |
| | alpha | 1405 | 165 | theta | 1427 | 156 | 0.2825 |
| | control | 1450 | 206 | NE | 1612 | 272 | 4.279* |
| | control | 1450 | 206 | theta | 1427 | 156 | 0.2773 |
| | NE | 1612 | 272 | theta | 1427 | 156 | 18.9926** |
| miss | alpha | 1580 | 257 | control | 1519 | 271 | 0.3652 |
| | alpha | 1580 | 257 | NE | 1771 | 344 | 3.3256* |
| | alpha | 1580 | 257 | theta | 1573 | 283 | 0.2472 |
| | control | 1519 | 271 | NE | 1771 | 344 | 19.4755** |
| | control | 1519 | 271 | theta | 1573 | 283 | 0.326 |
| | NE | 1771 | 344 | theta | 1573 | 283 | 3.3623* |
| CR | alpha | 1397 | 174 | control | 1417 | 236 | 0.265 |
| | alpha | 1397 | 174 | NE | 1609 | 296 | 31.4731** |
| | alpha | 1397 | 174 | theta | 1414 | 185 | 0.2642 |
| | control | 1417 | 236 | NE | 1609 | 296 | 6.9499* |
| | control | 1417 | 236 | theta | 1414 | 185 | 0.2463 |
| | NE | 1609 | 296 | theta | 1414 | 185 | 13.1874** |
| FA | alpha | 1669 | 222 | control | 1675 | 309 | 0.2469 |
| | alpha | 1669 | 222 | NE | 1885 | 310 | 37.8379** |
| | alpha | 1669 | 222 | theta | 1718 | 230 | 0.3493 |
| | control | 1675 | 309 | NE | 1885 | 310 | 9.4932* |
| | control | 1675 | 309 | theta | 1718 | 230 | 0.2961 |
| | NE | 1885 | 310 | theta | 1718 | 230 | 5.2933* |

*Note*. This table shows the results from the statistical analysis of reaction times based on entrainment group differences. CR = correct rejection, FA = false alarm, M = mean (arithmetic), SD = standard deviation, df = degrees of freedom

*a* The depicted Bayes factor values are equivalent to $BF_{10}$, estimating the evidence for the alternative hypothesis (statistical difference) relative to the null hypothesis (no difference). A Cauchy distribution of medium width was used as prior for each comparison.

* at least moderate evidence for $H_1$, ** at least strong evidence for $H_1$
