## Supplementary Information for "Disentangling the Functional Roles of Pre-Stimulus Oscillations in Crossmodal Associative Memory Formation via Sensory Entrainment"

### Supplementary material

**Jan Ostrowski[1†], Marike C. Maack[1†], Michael Rose[1*]**

1 *Department of Systems Neuroscience, University Medical Center Hamburg-Eppendorf, Hamburg, Germany;* **, **, **

† Equal contributions

*16-digit ORCID:*

*JO: 0000-0001-9928-272X*

*MM: 0000-0002-6960-4465*

*MR: 0000-0002-9789-7066*

**Supplementary 1. Contrasting pre-stimulus activity from theta and alpha groups to activity from the control and NE groups**

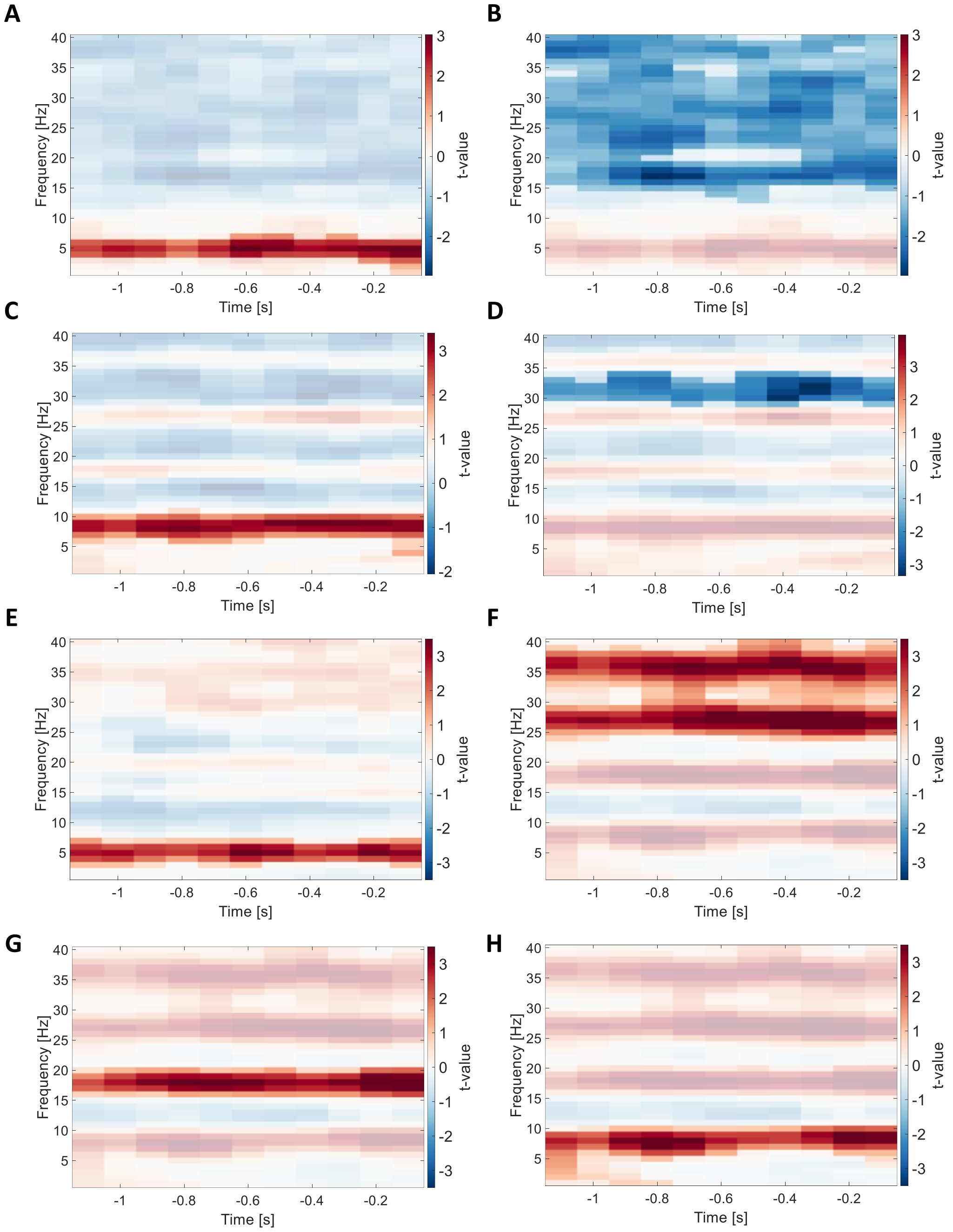


**Figure S1. Results of EEG data contrasts of entrainment groups with the control and NE groups.** The figure shows time-frequency plots depicting the results of the statistical comparison of relative change in pre-stimulus activity. **(A)** and **(B)** show the two significant clusters from the comparison of the theta group with the control group. **(C)** and **(D)** depict the two significant clusters resulting from contrasting the alpha group with the control group. **(E)** shows the positive cluster revealed by comparing activity from the theta group with the NE group. **(F) – (G)** depict the statistical results comparing the alpha group with the NE group. In all time-frequency plots, positive *t*-values signify greater relative change in the theta or alpha groups, respectively. Opaque data points show the extent of a statistically significant cluster ($p < .025$, corrected). Each subplot shows one distinct cluster and depicts the *t*-values averaged over the electrodes comprising the cluster.

Oscillatory power in the late entrainment period (-1.1 s to -0.1 s relative to stimulus onset) from the theta and alpha group was each contrasted with the activity from the NE group in the same time period. We used a cluster-based permutation approach to account for multiple comparisons, and two-tailed independent-samples *t*-tests on sample level. The frequency range for this analysis was set to 1 to 40 Hz, and all electrodes were included. Comparing the theta entrainment condition with the NE group, the analysis yielded one significant positive cluster in the frequency range of 3 to 7 Hz, spanning the whole analysis window ($p$ < .025, corrected). This suggests significantly increased oscillatory power in the envelope around 5 Hz for the theta group as compared to the NE group **(Figure S2E)**. Contrasting activity from the alpha group with the NE group revealed three distinct positive clusters, each spanning the whole analysis window. The clusters covered the frequency ranges of 24 to 40 Hz and 16 to 20 Hz. Importantly, the third cluster ranged from 1 to 10 Hz up until -0.8 s relative to stimulus onset and was centered one the 9 Hz envelope for the remaining part of the analysis time window (**Figure S2H**).

**Supplementary 2. Analysis of oscillatory activity in the stimulus presentation window during encoding**

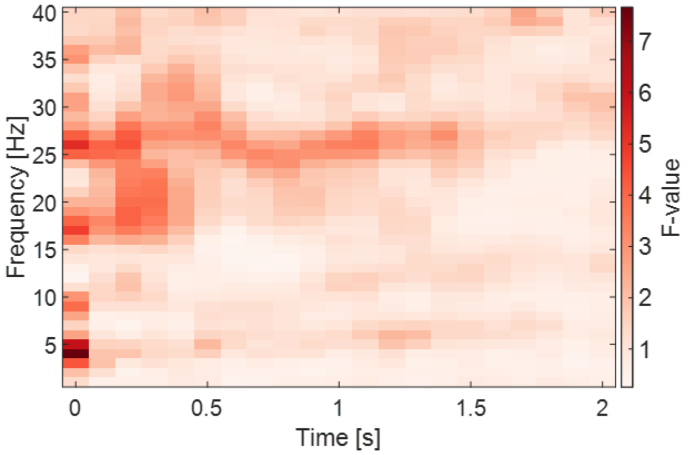


**Figure S2. Difference in post-stimulus activity between entrainment groups.** The figure shows a time-frequency plot of the stimulus presentation time window from the encoding task over a frequency range of 1 to 40 Hz. The color dimension displays the F-values from the independent-samples F-test.

In order to assess potential differences in oscillatory activity during stimulus presentation between groups, we compared oscillatory power from the post-stimulus interval (0 s to 2s relative to stimulus onset) among the entrainment groups (theta, alpha, and control) using an independent-samples *F*-test on sample level. Data was included for a frequency range of 1 to 40 Hz and all electrodes, and cluster-based permutation was used for multiple-comparison corrections. However, the analysis showed only a tendency for a significant cluster in the electrode-frequency-time space, suggesting that there are no significant differences in post-stimulus power between the entrainment groups ($p$ = .069, corrected).

**Supplementary 3. Analysis of categorization task performance during encoding**

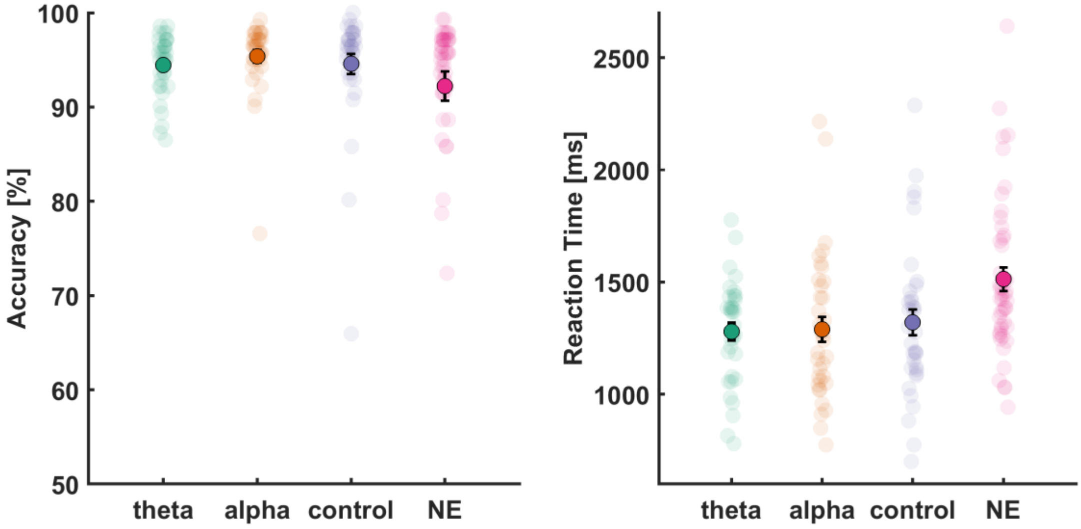


**Figure S3. Performance in the categorization task during encoding.** *Left*: Accuracy values for the categorization task during encoding for each group. *Right:* Average response time for the categorization task during encoding. Transparent data points mark individual task accuracy, and the black error bars signify the standard error of means.

In the categorization task from the encoding phase, participants showed high accuracy across all entrainment conditions, 94.45% (SD = 3.23) for theta, 95.38% (SD = 3.88) for alpha, and 94.57% (SD = 6.27) for control condition. Accuracy in the NE condition as an additional control was slightly lower at 92.21% (SD = 10.43). The overall accuracy across all four conditions was 94.02% (SD = 6.97). Using a Bayesian one-way ANOVA model yielded a Bayes factor of $BF_{10}$ = 0.381, suggesting moderate evidence in support of the null hypothesis of no significant differences among the groups. These results suggest that participants in the current study maintained high compliance with task demands throughout the experiment, which was essential for accurately assessing the subsequent impact of oscillatory activity on memory performance. Reaction times showed a similar pattern. Participants responded fastest in the entrainment conditions (theta: 1279.4 ms, SD = 230.3; alpha: 1288.9 ms, SD = 327.6; control: 1320.4 ms, SD = 343.7), with slower responses in the NE condition (1512.6 ms, SD = 355.3). The overall average reaction time was 1361.11 ms (SD = 333.1). However, a Bayesian one-way ANOVA indicated moderate evidence for the alternative hypothesis, $BF_{10}$ = 5.0737, indicating measurable differences among the groups. Individual group contrasts revealed that there was likely no difference in response times between the entrainment groups (theta vs control: $BF_{10}$ = 0.2849; alpha vs control: $BF_{10}$ = 0.2628; theta vs alpha: $BF_{10}$ = 0.248). However, the evidence suggests a moderate-to-strong effect for differences between the theta and alpha groups and the NE group (theta vs NE: $BF_{10}$ = 19.1684; alpha vs NE: $BF_{10}$ = 5.7022). The comparison between the control and the NE group yielded only weak evidence for significant difference, $BF_{10}$ = 2.3366.

**Supplementary 4.** ***Changes in performance over the course of the experiment***

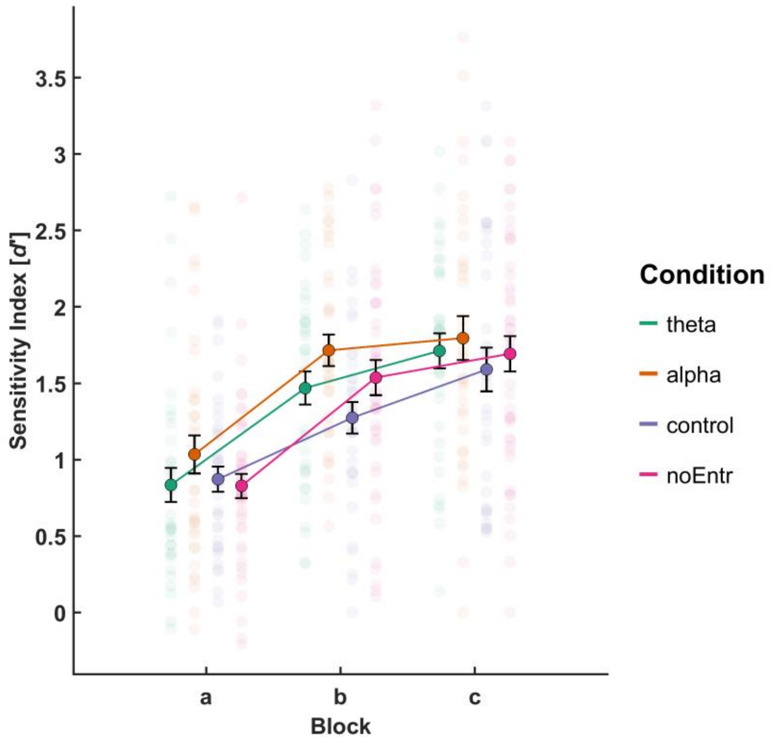


**Figure S4. Changes in sensitivity indices across experimental runs.** The plot depicts mean sensitivity indices (*d'*) over participants for every group and the three experimental runs. Every transparent data point marks the individual sensitivity index of one participant. Black error bars indicate the standard error of means.

* at least moderate evidence for $H_1$, ** at least strong evidence for $H_1$

**Supplementary 6. *Resting State***

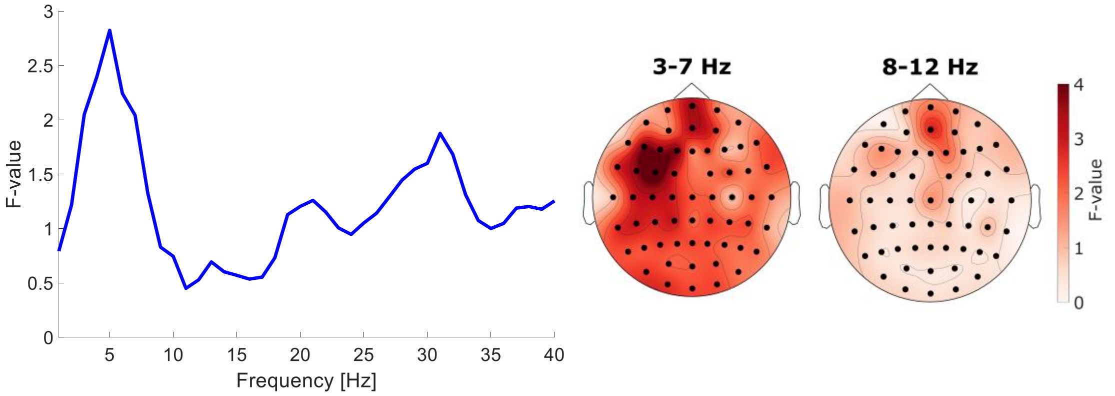


**Figure S5. Group contrast for differences in resting-state activity before and after the experiment.** *Left*: F-values for the analysis window of 1 to 40 Hz from a cluster-based permutation test with an independent-samples *F*-test on sample level assessing group differences in pre- and post-experiment resting-state discrepancies. . *Right*: Topographical distribution of *F*-values averaged across the 3–7 Hz (theta) and 8–12 Hz (alpha) bands. No significant clusters were observed in this analysis (p = .2972, corrected).

As some studies report lingering oscillatory effects due to entrainment procedures **[Kasten & Herrmann, 2022; Gallina et al., 2023)**, we explored differences in resting-state spectra that were recorded once before (RestPre) and once after the SME task (RestPost) to determine whether traces of the entrainment could be observed even after the experiment. For the analysis, we used power spectra in the frequency range of 1 to 40 Hz and submitted the data to a cluster-based permutation test with two-tailed paired-samples t-tests on the sample level. Note that this analysis was conducted separately for the theta group, alpha group, as well as the control group. For the theta group, one negative cluster was found in the alpha band (8 to 12 Hz), indicating increased power after the experiment (p < .025, corrected). Similarly, a negative cluster ranging from 7 to 18 Hz was observed for the comparison in the alpha group (p < .025, corrected), while the analysis in the control revealed a negative cluster in the alpha band (8 - 12 Hz, p < .025, corrected). Interestingly, the control group analysis yielded a second negative cluster in the beta band, ranging from 17 to 33 Hz (p < .025, corrected). As the effect in the alpha band and, to a certain degree, in the beta band was observed in all three groups, we were interested in whether the effect magnitude differed between the groups.

**Supplementary 7.** ***No differences in subjective perception of entrainment***

Survey items (translated from German into English):

*I1 (pleasentness)*: How pleasant did you find the flickering of the image?

*I2 (distraction)*: To what extent did you feel distracted by the flickering of the image while trying to remember the pairs?

*I3 (attention)*: How would you rate your level of attention during the task?

*I4 (fatigue)*: How exhausted do you feel at the moment?

Participants rated on a scale from 0 (*not at all*) to 5 (*very much*) in steps of 0.5.

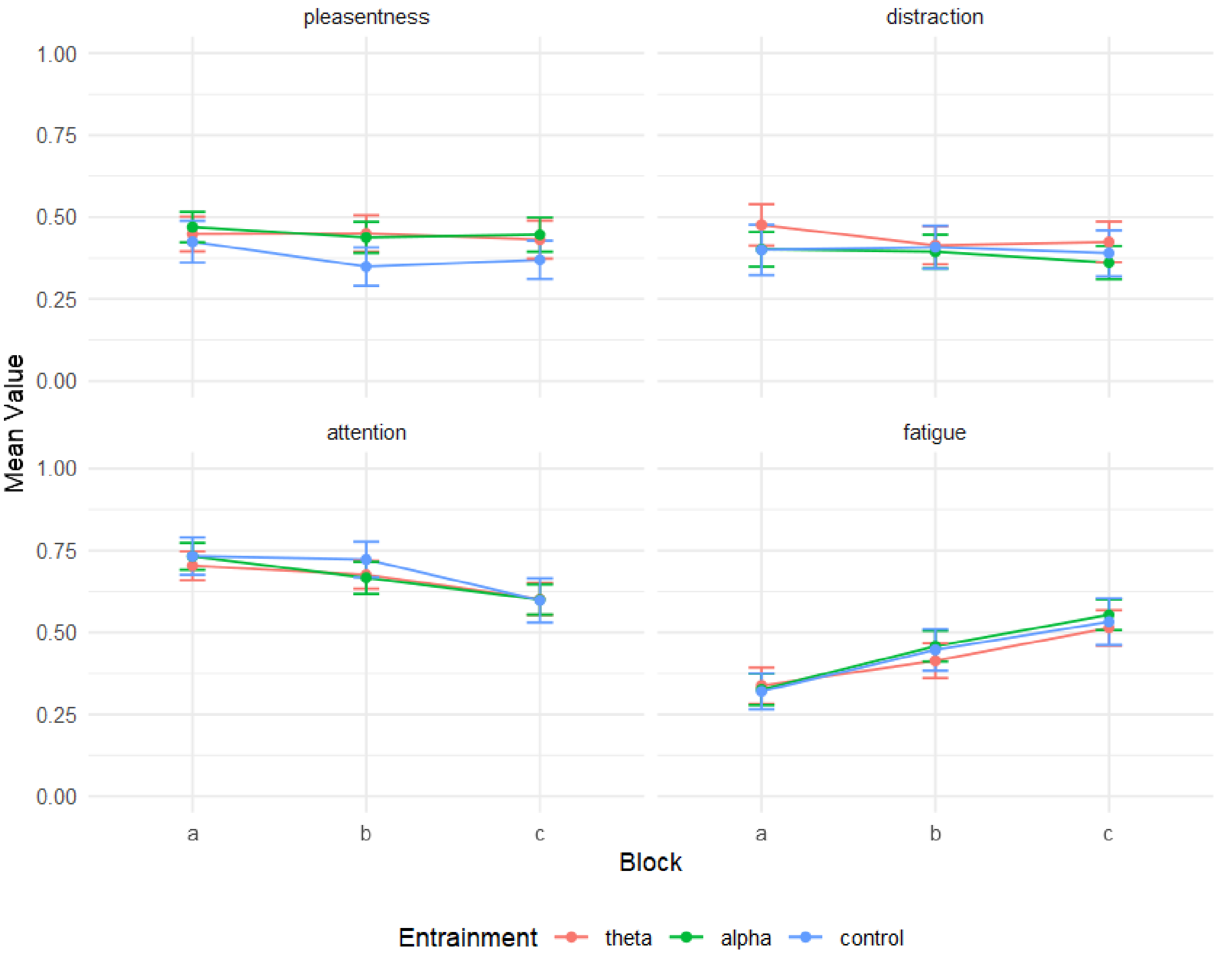


**Figure S6**. Subjective ratings of task-related experience across blocks and entrainment conditions. The data points depict group and block averages across participants. The error bars mark the standard error of means.

As individual perception qualities of images with oscillating luminance may vary, we investigated whether the subjective perception of the sensory stimulation might differ between the entrainment groups to control for salience effects. Participants received four survey items after each encoding phase, measuring the *pleasantness* and the *distracting qualities* of the entrainment procedure, as well as *attention* and *fatigue*. We conducted a Bayesian mixed-design ANOVA for every item, with a between-subjects factor *pre-stimulus condition* (theta, alpha, control) and a within-subjects factor *block* (A, B, C). Scores from every item did not significantly differ between levels of *pre-stimulus condition* ($BF_{pleasentness}$ = 0.3099, $BF_{distraction}$ = 0.3853 , $BF_{attention}$ = 0.1132 , $BF_{fatigue}$ = 0.1279 ), indicating that the type of entrainment procedure had no differential effect. However, the analysis revealed an effect of *block* for the variables *distraction, attention,* and *fatigue* ($BF_{distraction}$ = 7.8834, $BF_{attention}$ = $1.855 \times 10^9$, $BF_{fatigue}$ = $1.3415 \times 10^{15}$). No interactions of *pre-stimulus condition* and *block* were observed ($BF_{pleasentness}$ = 0.0732, $BF_{distraction}$ = 0.0578, $BF_{attention}$ = 0.195, $BF_{fatigue}$ = 0.0552. Participants felt less distracted by the entrainment in block C of the experiment (M = 0.382, SD = 0.224) than in block A (M = 0.45, SD = 0.243). Conversely, participants rated their level of attention in block C (M = 0.573, SD = 0.203) consistently lower than in block A (M = 0.694, SD = 0.186). This was accompanied by increased fatigue ratings in block C (M = 0.528, SD = 0.223) as compared to block A (M = 0.338, SD = 0.197). In sum, evidence from the survey data indicates that the entrainment procedures were received equally pleasant and distracting, suggesting no confound of the behavioral results due to subjective perception.
